# Elucidation of chalkophomycin biosynthesis reveals *N*-hydroxypyrrole-forming enzymes

**DOI:** 10.1101/2024.01.24.577118

**Authors:** Anne Marie Crooke, Anika K. Chand, Zheng Cui, Emily P. Balskus

## Abstract

Reactive functional groups, such as *N*-nitrosamines, impart unique bioactivities to the natural products in which they are found. Recent work has illuminated enzymatic *N*-nitrosation reactions in microbial natural product biosynthesis, motivating an interest in discovering additional metabolites constructed using such reactivity. Here, we use a genome mining approach to identify over 400 cryptic biosynthetic gene clusters (BGCs) encoding homologs of the *N*-nitrosating biosynthetic enzyme SznF, including the BGC for chalkophomycin, a Cu^II^-binding metabolite that contains a *C*-type diazeniumdiolate and *N*-hydroxypyrrole. Characterizing chalkophomycin biosynthetic enzymes reveals previously unknown enzymes responsible for *N*-hydroxypyrrole biosynthesis, including the first prolyl-*N*-hydroxylase, and a key step in assembly of the diazeniumdiolate-containing amino acid graminine. Discovery of this pathway enriches our understanding of the biosynthetic logic employed in constructing unusual heteroatom-heteroatom bond-containing functional groups, enabling future efforts in natural product discovery and biocatalysis.

## INTRODUCTION

Many natural product bioactivities are a direct consequence of incorporating reactive functional groups. *N*-nitroso groups, for example, are precursors to DNA alkylating agents and generate reactive NO radicals.^1–4^ Recently, diazeniumdiolates (*N*-hydroxy-*N*-nitrosamines) have been identified as a bidentate ligand in multiple metallophores.^5–7^ This plethora of functions has increased interest in *N*-nitroso natural products and in discovering biosynthetic enzymes that synthesize this and similarly reactive functional groups containing heteroatom-heteroatom bonds.^8–12^

The first dedicated *N*-nitrosating enzyme to be biochemically characterized, SznF, was identified in the biosynthesis of the FDA-approved cancer chemotherapeutic streptozotocin.^13,14^ This multidomain metalloenzyme catalyzes the oxidative rearrangement of a guanidine group to the *N*-nitrosourea pharmacophore of streptozotocin (Figure 1A, Figure S1). The diiron-binding heme oxygenase-like diiron oxidase and oxygenase (HDO) domain of SznF converts L-*N*^ω^-methylarginine to L-*N*^δ^-hydroxy-*N*^ω^-hydroxy-*N*^ω’^-methylarginine, and its non-heme mononuclear iron-containing cupin domain catalyzes a synthetically and biochemically unprecedented intramolecular rearrangement of this intermediate to afford the *N*-nitrosourea product.^15,16^ Prior to the discovery of SznF, characterized approaches for *N*-nitrosation in biology invoked non-enzymatic nitrosation of amines using nitrite; however, the evolution of a dedicated *N*-nitrosating enzyme suggests important biological roles for this functional group and raises the questions about the distribution of this enzymatic chemistry.

**Figure 1.**
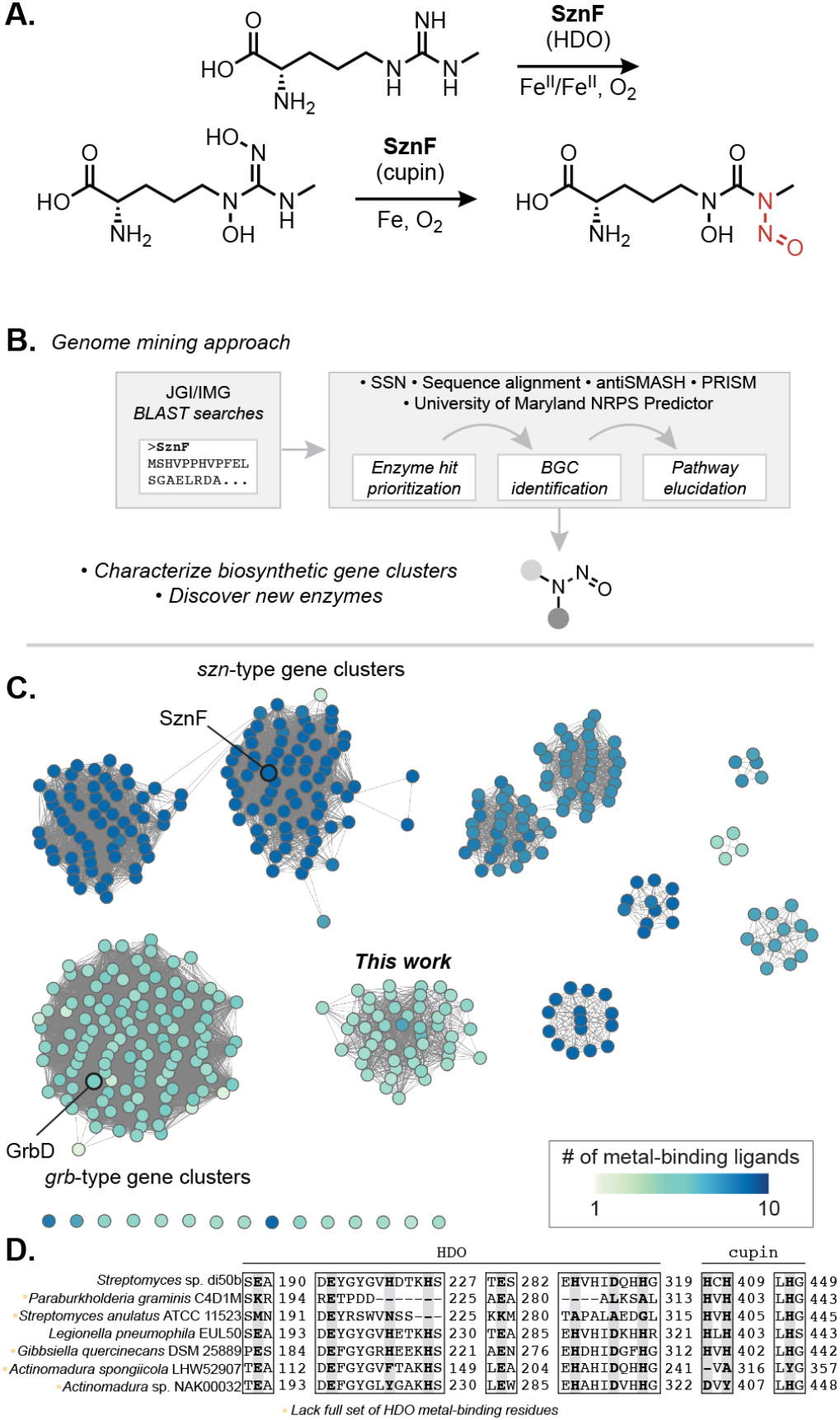
Genome mining reveals numerous uncharacterized biosynthetic pathways that use putative *N*-nitrosating enzymes. (A) SznF is a non-heme iron-dependent oxygenase that catalyzes guanidine *N*-oxygenation and *N*-nitrosation in streptozotocin biosynthesis. Leveraging an understanding of SznF-catalyzed chemistry may inform the discovery of new *N*-nitrosating enzymes. (B) Workflow for identification and characterization of biosynthetic gene clusters (BGCs) that make putative *N*-nitroso natural products. (C) Sequence similarity network (SSN) of 426 unique protein sequences identified through BLAST search in the JGI/IMG database of sequenced microbial genomes (E-value = 1e– 110). Multiple Sequence Comparison by Log-Expectation (MUSCLE) alignment was used to determine conservation of the metal-binding residues of SznF’s HDO (7 residues) and cupin (3 residues) domains. The node highlighting SznF represents the protein sequences from *Streptomyces* sp. di50b and *S*. sp. di188 (94% amino acid ID to query SznF sequence from *S. achromogenes* var. *streptozoticus* NRRL 2697). (D) Select examples from the MUSCLE alignment of all protein sequences derived from the seven largest clusters in the SSN. The position of metal binding residues of the heme oxygenase-like diiron oxidase and oxygenase (HDO) domain and the cupin domain in SznF’s sequence are highlighted (bolded, grey) to convey the diversity in ligand conservation across BLAST hits.

Genome mining has been a successful strategy for discovering novel biosynthetic gene clusters (BGCs), including several pathways that produce N–N bond-containing natural products. *N*-nitroso metabolite-producing BGCs discovered using this strategy include the gramibactin, megapolibactin, plantaribactin, and tistrellabactin BGCs, all of which produce diazeniumdiolate (*N*-hydroxy-*N*-nitroso)-containing metallophores.^5,6,17^ In each case, a gene encoding an SznF homolog that is believed to carry out *N*-nitrosation has been identified in the corresponding BGC; however, these enzymes have not been biochemically characterized. Previous work from our group demonstrated the wide distribution of SznF homologs in phylogenetically diverse bacteria.^13^ However, compared to the number of unique BGCs predicted to encode an *N*-nitrosating enzyme, very few *N*-nitroso-containing natural products have been isolated.

Here, we use genome mining to identify additional *N*-nitrosating enzyme-encoding BGCs, including the BGC that produces chalkophomycin, a Cu^II^-binding metallophore that features the *C*-type diazeniumdiolate-containing amino acid graminine and a rare *N*-hydroxypyrrole heterocycle.^7^ We elucidate the biosynthetic origins of these unusual functional groups using stable isotope feeding experiments and further probe the enzymes responsible for their construction using *in vitro* biochemical characterization. We unexpectedly find that a heme-dependent enzyme participates in *N*-nitroso biosynthesis by generating L-dihydroxyarginine, akin to the role of SznF’s HDO domain, providing the first biochemical insights into the origins of graminine. We also characterize the enzymes responsible for biosynthesizing *N*-hydroxypyrrole from L-proline, uncovering a biosynthetic logic distinct from that used in the assembly of other functionalized pyrroles. Identifying the genetic and biochemical basis for *N*-hydroxypyrrole biosynthesis solves a longstanding mystery in the field, as the enzymes used to construct this structural motif have never been reported, despite its presence in natural products being noted for decades. This knowledge has enabled us to identify additional cryptic BGCs that likely produce *N*hydroxypyrroles, suggesting this heterocycle may play important roles in many more uncharacterized natural products.

## RESULTS

### Genome mining identifies the chalkophomycin biosynthetic gene cluster

To identify putative *N-*nitrosating enzymes encoded in uncharacterized BGCs, we performed iterative Basic Local Alignment Search Tool (BLAST) searches of the Joint Genome Institute/Integrated Microbial Genomes (JGI/IMG) database using the full amino acid sequence of SznF (Figure 1B).^18^ We classified these proteins into putative isofunctional groups using a Sequence Similarity Network (SSN) generated using the EFI-Enzyme Similarity Tool (Figure 1C).^19^ A large cluster of 118 sequences in the SSN contained enzymes previously implicated in diazeniumdiolate biosynthesis (GrbD, MegD, PlbJ, MobE), although they have not yet been biochemically characterized. A second SSN cluster included SznF and 135 other proteins located in *szn*-type BGCs.

Finally, a third prominent cluster in the SSN contained 48 uncharacterized proteins not yet known to perform *N*-nitrosation chemistry. The genes encoding for these proteins are found in a variety of unique genomic contexts. Unlike *sznF*, but like *grbD*, these genes co-localize with genes encoding nonribosomal peptide synthetase (NRPS) and polyketide synthase (PKS) machinery. We therefore predicted these enzymes biosynthesize distinct nonribosomal peptide and polyketide natural products featuring an *N*-nitroso functional group.

We next sought to assess the chemical capabilities of the SznF homologs in the SSN. A multiple sequence alignment of all 426 identified proteins was performed to examine conservation of the metal-binding residues in the HDO and cupin domains of SznF (Figure 1D). Intriguingly, among the sequences in the newly defined cluster of NRPS- and PKS-associated enzymes, all but one lack many of the iron-binding residues of the HDO domain but retain the three histidines of SznF’s N–N bond-forming cupin domain. This observation suggests these SznF homologs may lack *N*-oxygenating activity but might still catalyze a N–N bond formation.

To further explore the biosynthetic roles of these enzymes, we selected the BGC from *Streptomyces anulatus* ATCC 11523 for further investigation due to the strain’s commercial availability and well-annotated genome (JGI/IMG Genome ID: 2873289101). An identical BGC is also encoded in the genome of *Lentzea flaviverrucosa* DSM 44664. Comparing the genomes of these two strains revealed the conservation of 17 consecutive genes that compose the BGC of interest (Figure 2A). Using a combination of gene annotations, conserved domain databases from NCBI and InterPro, and NRPS-PKS analysis databases (antiSMASH, University of Maryland NRPS Predictor, and PRISM)^20–22^ we generated a predicted structure for the encoded natural product to inform metabolite identification efforts (Table S1, S2). Although there was ambiguity surrounding substrate predictions for many of the biosynthetic enzymes, we deduced that in addition to an *N*-nitroso group, this metabolite likely contained features commonly found in metallophores, including a catechol or phenol as well as a thiazoline heterocycle (Table S2).

**Figure 2.**
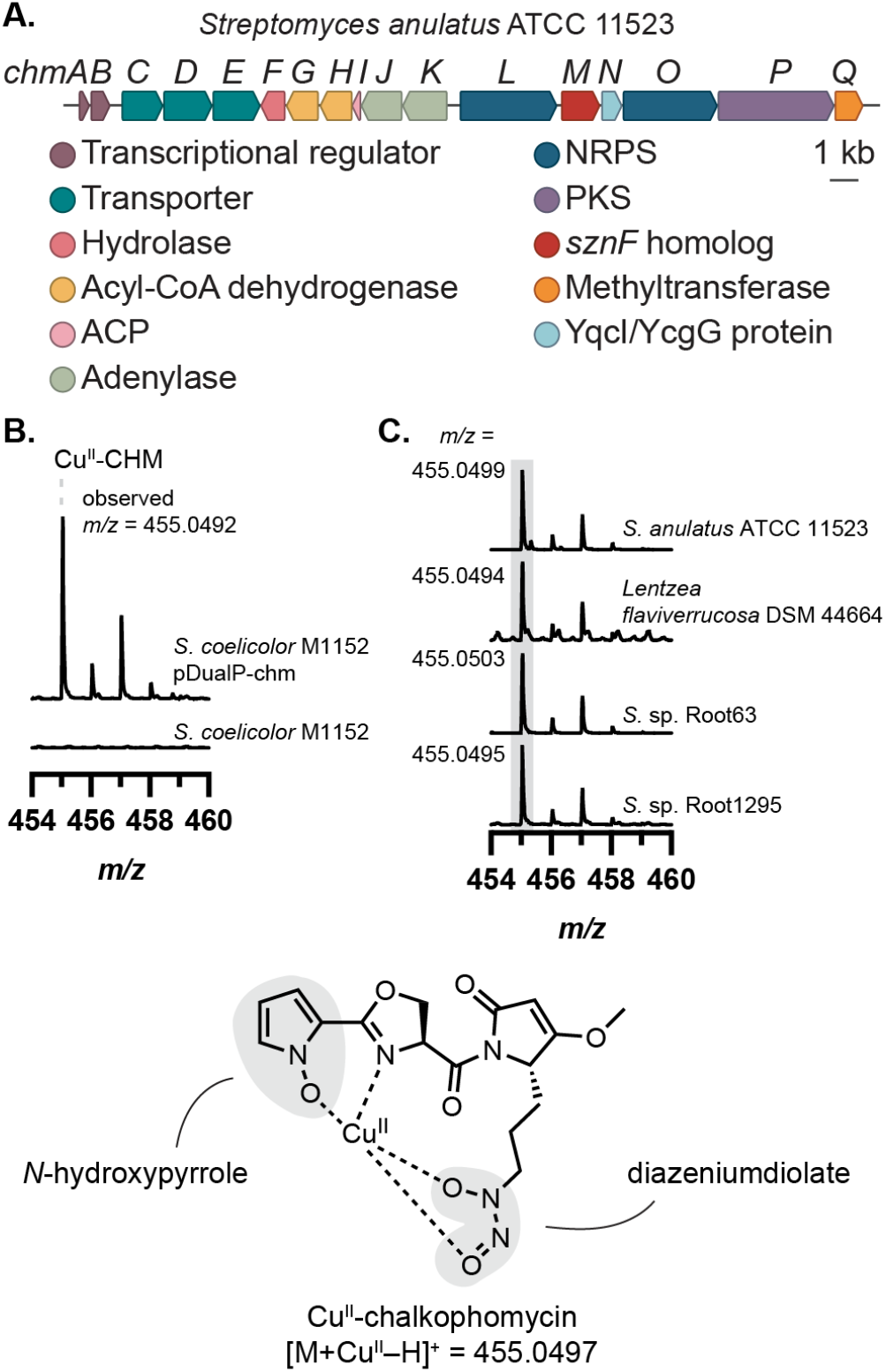
The *chm* gene cluster produces the metallophore chalkophomycin. (A) The *sznF* homolog *chmM* is encoded within a biosynthetic gene cluster (BGC) in the *Streptomyces anulatus* ATCC 11523 genome. (B) Heterologous expression of the *chm* gene cluster in *Streptomyces coelicolor* M1152 reveals chalkophomycin production. (C) The same mass feature, corresponding to chalkophomycin, can be observed in metabolomic samples of native *chm*-encoding organisms with the same *m/z* ratio and diagnostic Cu isotopic distribution pattern. Mass spectrum intensity was normalized to the height of 455 *m/z* ion in each sample.

We recognized a striking resemblance between elements of our predicted structure and chalkophomycin, a Cu(II)-binding natural product from *Streptomyces* sp. CB00271 that was reported shortly after we had completed our genome mining effort.^7^ Notably, chalkophomycin contains an *N*-hydroxypyrrole in lieu of the predicted catechol or phenol, where the N–OH serves as a metal-binding ligand. Chalkophomycin also has an oxazoline in place of the predicted thiazoline. Finally, the presence of a diazeniumdiolate in this natural product is consistent with a biosynthetic pathway that involves an SznF homolog. A BLAST search of the *S*. sp. CB00271 genome using SznF revealed a hit that shared 97% amino acid identity (aa ID) to the *S. anulatus* ATCC 11523 homolog, ChmM, encoded within an identical BGC. This provided strong support for the involvement of this cryptic gene cluster (which we termed the *chm* gene cluster) in chalkophomycin production.

To confirm this assignment, we heterologously expressed the *chm* BGC from *S. anulatus* ATCC 11523 in *Streptomyces coelicolor* M1152 (Figure 2B). *S. coelicolor* M1152 harboring the *chm* gene cluster (*S. coelicolor* pDualP-chm), but not the wild-type strain, produced chalkophomycin in R2B medium supplemented with 100 mg/L Cu(II)SO_4_ • 5H_2_O, confirmed by liquid chromatography–mass spectrometry (LC– MS) with the exact mass and Cu-specific isotopic distribution pattern expected for this metabolite. When *S. anulatus* ATCC 11523 was cultured in the medium used for chalkophomycin isolation (M2 medium), chalkophomycin was detected via LC–MS. Additionally, *L. flaviverrucosa* DSM 44664 and several other strains possessing the *chm* BGC also produced chalkophomycin under the R2B + Cu(II) growth conditions (Figure 2C, Figure S2–3). Apo-chalkophomycin was identified by LC–MS, and the MS/MS fragmentation pattern matches reported data (Figure S4).^7^ Therefore, heterologous expression of the *chm* BGC and metabolite profiling of native encoders unambiguously verified the link between the *chm* gene cluster and chalkophomycin.

We next sought to formulate an initial hypothesis for chalkophomycin biosynthesis and assign roles for individual Chm biosynthetic enzymes (Figure 3). We proposed a convergent biosynthesis that employs thiotemplated-formation of *N*-hydroxypyrrole and L-graminine, the diazeniumdiolate-containing non-proteinogenic amino acid. In this report, we document the enzymatic transformations that afford the *N*-hydroxypyrrole building block and intermediates in L-graminine synthesis. We hypothesized that the *N*-hydroxypyrrole is hydrolyzed from ChmI and re-introduced as the starter unit of the NRPS assembly line beginning with ChmL, after which the remaining amino acid substrates are incorporated into the growing natural product structure. ChmP adds a unit of malonyl-CoA, the last substrate required to produce the core chalkophomycin scaffold; however, there is no clear thioesterase (TE) domain in ChmP. The cyclization could occur via a non-enzymatic reaction or a TE located elsewhere in the genome. Cyclization by an unknown mechanism yields the dehydrolactam, and a final *O-*methylation would afford the final natural product.

**Figure 3.**
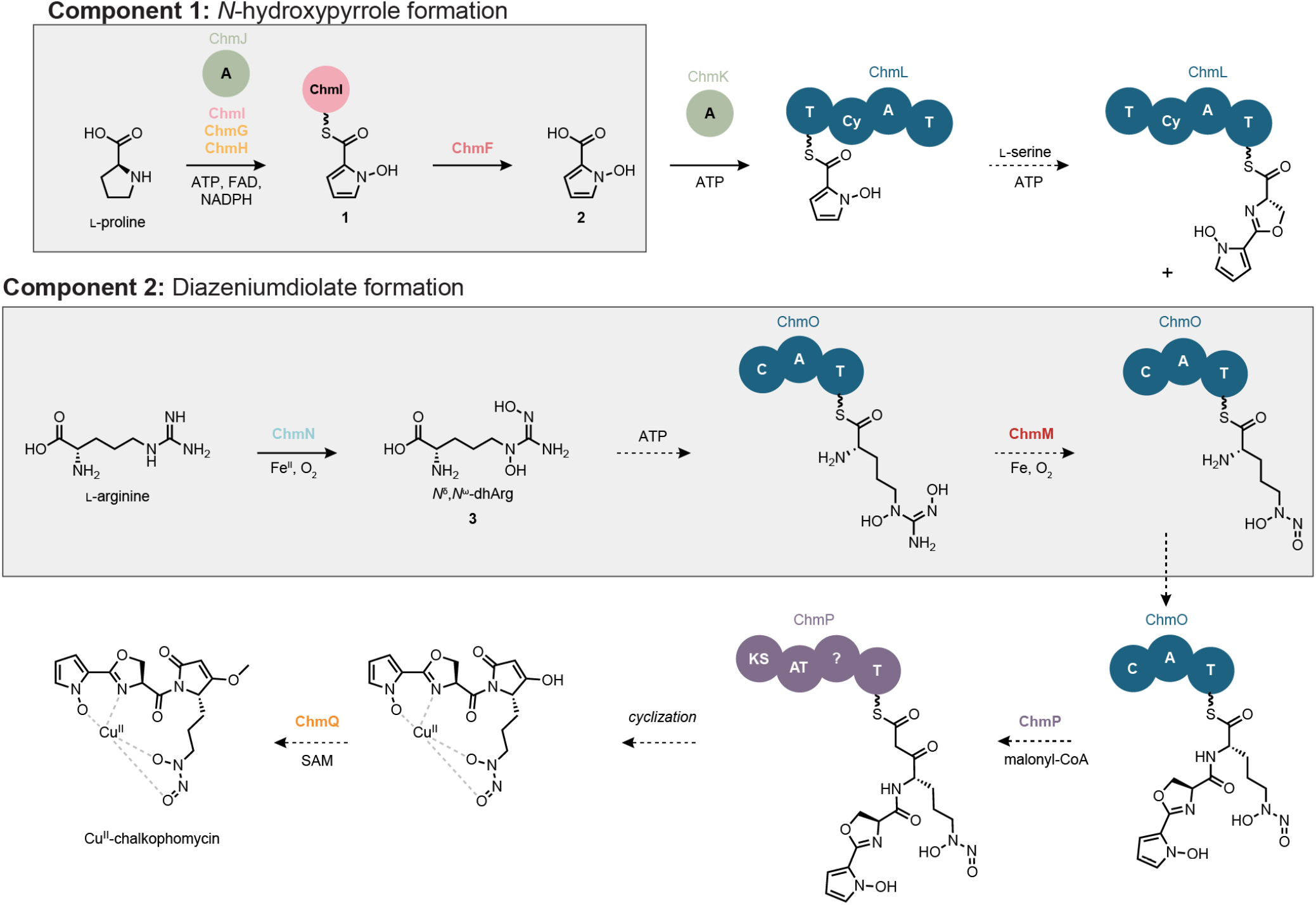
Hypothesis for chalkophomycin biosynthesis. NRPS and PKS domain abbreviations: T = thiolation, Cy = cyclization, C = condensation, A = adenylation, KS = ketosynthase, AT = acyltransferase. Cofactor abbreviations: ATP = adenosine triphosphate, FAD = flavin adenine dinucleotide, NADPH = nicotinamide adenine dinucleotide phosphate, SAM = *S*-adenosylmethionine.

### Stable isotope feeding experiments confirm biosynthetic origins of reactive functional groups

To determine the biosynthetic precursors of the *N*-hydroxypyrrole and diazeniumdiolate functional groups, we performed stable isotope feeding experiments. Noting the peptidic nature of chalkophomycin, we hypothesized that *N*-hydroxypyrrole would derive from L-proline through a series of oxidations, analogous to one established pathway for pyrrole biosynthesis.^23^ We proposed the L-graminine residue could either arise from L-arginine via an SznF-type rearrangement or from an intermolecular *N*-nitrosation reaction between nitrite and L-ornithine. To test these proposals, ^15^N-L-proline, ^15^N_4_ ^13^C_6_ -L-arginine, ^15^N_2_-L-ornithine, or ^15^N-sodium nitrite was added to cultures of *S*. sp. Root63 and culture supernatants were analyzed by LC–MS/MS. We chose this strain due to its consistently robust production of chalkophomycin (LC–MS peak areas ∼1.2–3.2 x 10^7^). Significant mass enrichment of chalkophomycin was only observed for the ^15^N-L-proline and ^15^N_4_ ^13^C_6_ -L-arginine fed cultures (Figure 4A). Moreover, in ^15^N-L-proline-fed cultures the label was isolated to fragments containing the *N*-hydroxypyrrole, and in arginine-fed cultures mass enrichment of the diagnostic NO-loss localized the arginine-derived labels to the diazeniumdiolate group (Figure 4B, Figure S5). Consistent with our findings, while our studies were ongoing, efforts to characterize the biosynthetic origin of L-graminine in gramibactin and tistrellabactin biosynthesis also revealed L-arginine as the source of the diazeniumdiolate.^6,24^

**Figure 4.**
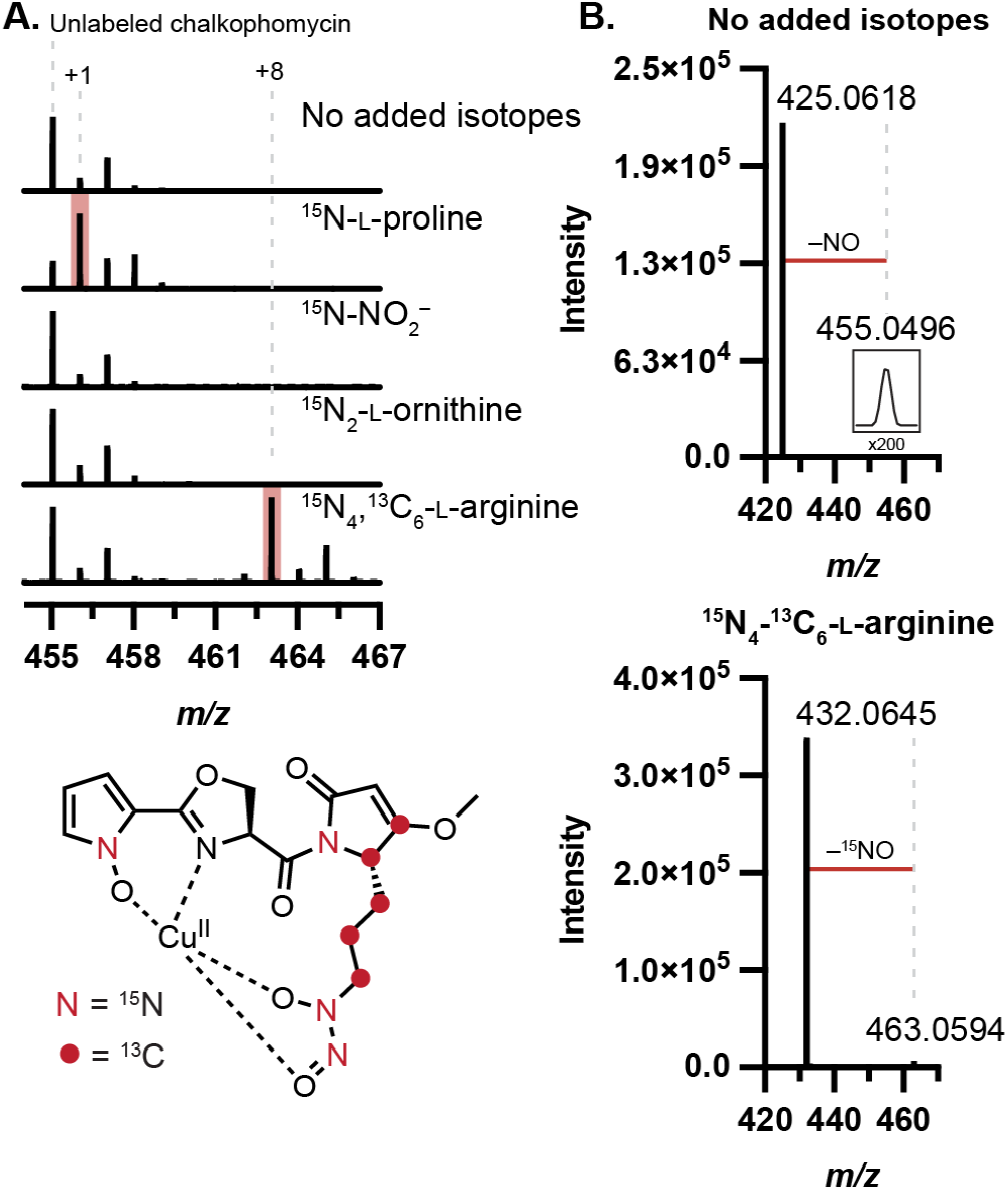
Stable isotope labeling determines precursors for chalkophomycin biosynthesis. (A) ^13^C and ^15^N are incorporated into chalkophomycin from L-proline and L-arginine by metabolomic analysis of *S*. sp. Root63 cultures supplemented with isotopically-enriched substrate. Percent enrichment for each substrate (Mean ± SD%): ^15^N-L-proline (64.93 ± 9.52%), ^15^N-sodium nitrite (1.88 ± 0.08%), ^15^N_2_-L-ornithine (11.65 ± 4.16%), ^15^N_4_ ^13^C_6_ -L-arginine (49.03 ± 3.58%). (B) LC–MS/MS of chalkophomycin results in ^15^NO fragment loss when ^15^N_4_,^13^C_6_-L-arginine is incorporated, localizing arginine-derived atoms to the diazeniumdiolate.

### *N*-hydroxypyrrole biosynthesis requires two flavin-dependent enzymes

With key chalkophomycin biosynthetic precursors identified, we next investigated the activities of enzymes responsible for constructing the unusual *N*-hydroxypyrrole heterocycle. *N*-hydroxypyrroles are rarely found in natural products but have been observed previously in hormaomycin, glycerinopyrin, pyranonigrins B and C, and surugapyrroles A and B.^25–30^ Chalkophomycin is the first example of this functional group serving as a metal-binding ligand. Additionally, pyrroles are a widespread motif in pharmaceuticals.^31,32^ The *N*-hydroxypyrrole heterocycle likely has several distinct biosynthetic origins, including from L-leucine and a linear PKS product (Figure S7).^33–36^ However, in all cases the enzymatic chemistry involved in *N*-hydroxypyrrole formation has eluded characterization.

Analysis of the *chm* BGC revealed four genes encoding likely candidate *N*-hydroxypyrrole-forming enzymes (*chmGHIJ*). A closely related set of genes is also found in the hormaomycin BGC (*hrmKLMN*); however, these genes and their encoded enzymes have never been investigated.^37^ Three of these genes share annotations with several previously characterized enzymes required for thiotemplated pyrrole biosynthesis: an acyl carrier protein (ACP), a proline-specific adenylating enzyme, and an FAD-dependent acyl-CoA dehydrogenase (ACAD).^23,38^ The adenylating enzyme uses ATP to activate L-proline via a prolyl-AMP intermediate that is transferred onto the thiol of the phosphopantetheine (ppant) arm of the ACP. The ACAD then catalyzes a 4-electron oxidation of the pyrrolidine ring of the prolyl thioester to the corresponding pyrrole prior to further elaboration of the pyrrole scaffold.^38,39^ The *chm* gene cluster encodes an ACP (ChmI) and an adenylase (ChmJ); however, it encodes two putative ACADs (ChmG and ChmH). We envisioned the second ACAD might perform *N*-oxygenation post-pyrrole formation.^40^

To elucidate the biosynthetic logic of *N*-hydroxypyrrole formation, we reconstituted its biosynthesis *in vitro* (Figure 5A). ChmI and ChmJ were recombinantly expressed in *E. coli* and their activity toward Lproline examined (see Supporting Information for further details). When *holo*-ChmI and ChmJ were incubated with ATP and L-proline, we observed L-proline loading onto the ppant arm of ChmI via LC–MS (**4**), which was further confirmed by targeted MS/MS to release a diagnostic ppant-proline fragment (Figure 5C, E). When ChmJ, ATP, or Lproline were omitted from assay mixtures, prolyl-ChmI (**4**) was not observed (Figure S8), indicating that ChmJ activates L-proline for loading onto the ppant arm of ChmI.

**Figure 5.**
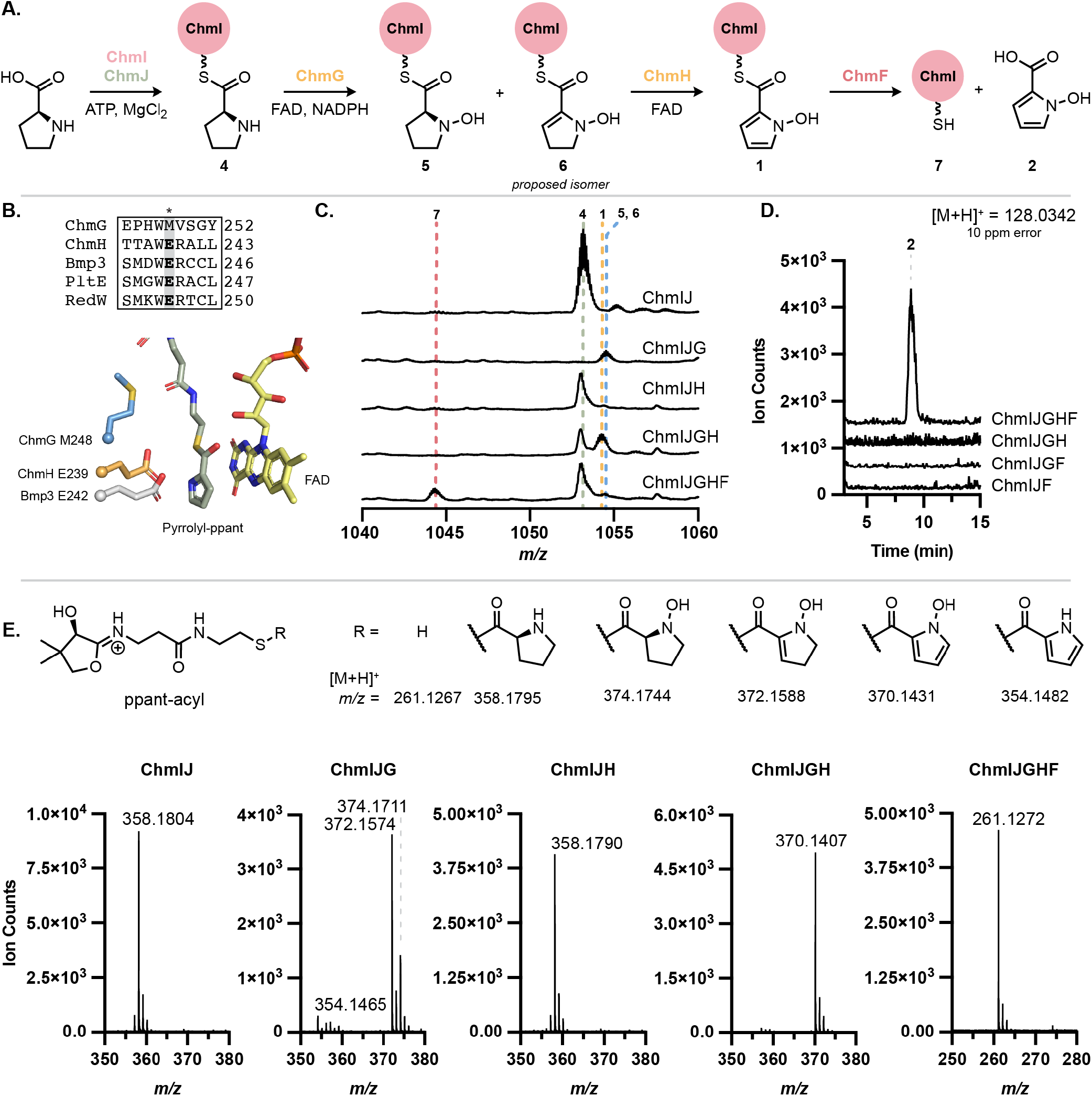
*N-*oxygenation precedes pyrrole formation in the biosynthesis of *N-*hydroxypyrrole. (A) L-Proline is converted to *N*-hydroxypyrrole-2carboxylic acid by ChmIJGHF. Note the C2–C3 double bond in **6** is proposed based on literature precedent but has not been experimentally confirmed. (B) MUSCLE alignment of ChmG and ChmH with characterized pyrrole-forming enzymes Bmp3, PltE, and RedW shows that only ChmH retains the catalytic glutamate required for pyrrolidine dehydrogenation. AlphaFold2 models of ChmH and ChmG superimposed with Bmp3 crystal structure (PDB: 6CXT) shows spatial alignment of the glutamate and methionine residues, respectively. Pyrrolyl-ppant (green) and FAD (yellow) are from the Bmp3 structure. (C) Whole protein LC–MS activity assay demonstrates that *Lfl*ChmG and *Lfl*ChmH are required for modification of ChmI-proline substrate. (D) Free *N*-hydroxypyrrole-2-carboxylic acid is hydrolyzed from ChmI in the presence of ChmF, as observed by LC–MS. (E) LC–MS/MS of whole protein mass features in (C) confirm proline modification from observation of the ppant-derived mass features.

We next examined the candidate enzymes for L-proline oxidation. Both ChmG and ChmH are annotated as ACADs. ACADs, including RedW, PltE, and Bmp3, catalyze the 4-electron oxidation of carrier protein-bound L-proline to the corresponding pyrrole.^23,38,39^ Although ChmG and ChmH are most similar to RedW, they share only 24% and 31% aa ID with this protein, respectively. As such, it was not evident which of the two proteins may perform dehydrogenation and whether there may be differences between this system and characterized pyrrole-forming pathways. Moreover, ChmG and ChmH share only 22% aa ID to each other, suggesting that they might perform distinct chemical transformations.

Multiple sequence alignments revealed that ChmH has the conserved, catalytic glutamate residue essential for pyrrole formation in AnaB and Bmp3 (Figure 5B, Figure S9).^41,42^ This catalytic base is proposed to initiate substrate oxidation by deprotonation of C2–H to form a prolyl thioester enolate, facilitating subsequent hydride transfer to FAD. Although the exact position of hydride transfer (C3 or N1) has not been confirmed experimentally, QM/MM studies of the Bmp1–Bmp3 complex suggest hydride transfer from C3 as most favorable.^43^ Unlike the characterized proline oxidases, ChmG has a methionine at this position, suggesting it might catalyze a distinct reaction. Searches with the DALI server using an AlphaFold2-generated structure of ChmG revealed structural similarity to the flavin-dependent amino sugar *N*-oxygenase KijD3 (24% aa ID), suggesting ChmG might possess *N*-oxygenase activity (Table S3),^44–47^. This was not evident from ChmG’s sequence or gene annotation, making it unique in the ACAD superfamily for its proposed *N*-oxygenase activity towards a proline-derived thioester. Combined with a phylogenetic analysis of the broader “Acyl-CoA dehydrogenase” enzyme family (IPR006091) (Figure S10), these data led us to hypothesize that ChmH catalyzes the 4-electron oxidation of proline prior to ChmG-catalyzed *N*-oxygenation. This proposed logic would resemble other biosyntheses of functionalized pyrroles.

To test this proposal, we biochemically characterized ChmH and ChmG *in vitro*. After encountering significant difficulties expressing the enzymes from *S. anulatus*, we successfully expressed homologs from *L. flaviverrucosa* DSM 44664 (*Lfl*ChmG and *Lfl*ChmH) in *E. coli*. We first tested if either *Lfl*ChmG or *Lfl*ChmH could oxidize ChmI-tethered prolyl thioester **4** (Figure 5C, E). Notably, no activity towards **4** was observed when *Lfl*ChmH was added alone (Figure S11). By contrast, when *Lfl*ChmG alone was added to assay mixtures containing *holo*-ChmI, ChmJ, proline, and ATP, **4** was converted primarily to a species consistent with ChmI-*N*-hydroxydehydroproline (**6**). Control experiments omitting NADPH or adding a flavin reductase to the reaction mixture resulted in varied product ratios of **5** and **6**, with both being formed, but the more highly oxidized product, **6**, was more abundant under conditions with higher FADH_2_ concentrations (Figure S12A,B). To control for the role reactive oxygen species (ROS) may play in this reaction, superoxide dismutase (100 U/mL) and catalase (35 U/mL) were added to the reaction mixtures. Both **5** and **6** were formed when only *Lfl*ChmG was added, but the product distribution favored **5**, suggesting ROS may be accelerating the conversion of **5** to **6 (**Figure S12C). **1** was still formed with both *Lfl*ChmG and *Lfl*ChmH, confirming that dehydrogenation is enzyme-catalyzed (Figure S12D). Addition of *Lfl*ChmH following pre-accumulation of **5** and **6** results in formation of **1** (Figure S13). These results indicate that *N*-oxygenation occurs prior to dehydrogenation in the biosynthesis of **1** and that both **5** and **6** may be on-pathway intermediates.

Collectively, our results demonstrate that thiotemplated *N*-hydroxypyrrole biosynthesis uses two distinct flavin-dependent enzymes. ChmG first catalyzes *N*-oxygenation of ChmI-proline, followed by either a 2or 4-electron oxidation by ChmH to afford the final ChmI-bound *N*hydroxypyrrole (**1**). This order of events has never been previously observed for thiotemplated biosynthesis of functionalized pyrroles, which has exclusively involved pyrrole formation prior to further transformations, such as halogenation.^48^ Additionally, ChmG is first example of a prolyl *N*-oxygenase and the first non-assembly line-associated flavin-dependent *N*-oxygenase to act on an aminoacyl-ACP substrate, broadening our knowledge of these oxygenases.

### ChmF hydrolyzes *N*-hydroxypyrrole from ChmI

Located nearby *chmGHIJ* in the *chm* BGC is *chmF*, a gene encoding a predicted α,β-hydrolase. This BGC also encodes a putative adenylate-forming enzyme (ChmK) that is predicted to accept 2,3-dihydroxybenzoic acid or salicylic acid (Table S2). We hypothesized that ChmF hydrolyzes the *N*-hydroxypyrrole from ChmI and that ChmK adenylates the resulting *N*-hydroxypyrrole-2-carboxylic acid (**2**) for loading onto the NRPS ChmL. Alternatively, this hydrolase may remove incorrectly loaded substrates from a carrier protein. To test the proposal that ChmF hydrolyzes the *N*-hydroxypyrrole thioester, we added ChmF to an assay mixture containing ChmIJGH and all necessary cofactors. We found that ChmF hydrolyzed *N*-hydroxypyrrole from ChmI (Figure 5C–E). Both the ChmI-ppant (**7**) and free *N*-hydroxypyrrole-2-carboxylic acid (**2**) products were observed by LC–MS only when ChmF was included in the reaction. ChmF appears to specifically recognize the *N*-hydroxypyrrole substrate, as little or no activity was observed towards **4** or the ChmG reaction products, **5** and **6** (Figure S14). Although it may seem inefficient to generate free *N*-hydroxypyrrole for reloading onto the assembly line, perhaps this enables incorporation of *N*-hydroxypyrrole into other additional biosynthetic pathways. When ChmK and ChmL were added to this reaction, we observed loading of *N*-hydroxypyrrole2-carboxylic acid (**2**) onto the first thiolation domain of ChmL (Figure S15). This not only confirms that ChmF hydrolyzes *N*-hydroxypyrrole, but it identifies ChmK as an adenylase for *N*-hydroxypyrroles. This logic also differs from characterized biosynthetic pathways that generate pyrroles from L-proline, where the pyrrole is assembled on a carrier protein and transferred to the subsequent NRPS or PKS without hydrolysis.^49^

### Bioinformatic analysis reveals many *N*-hydroxypyrroleencoding BGCs

With an understanding of *N*-hydroxypyrrole biosynthesis, we next examined the extent to which additional BGCs may produce this rare, functionalized heterocycle. Using prettyClusters, a set of tools that facilitates the analysis and visualization of genomic neighborhoods for a gene of interest, we identified 4 distinct classes of BGCs from 97 different organisms that encode homologs of the *N*-hydroxypyrrole forming enzymes (Figure 6, Figure S16).^50^ Many of these BGCs lack genes encoding obvious *N*-nitrosating enzymes. Notably, BGCs encoding this biosynthetic machinery are found in multiple human and plant pathogens such as *Nocardia nova* and *Pseudomonas fluorescens*. This highlights the existence of additional, undiscovered *N*-hydroxypyrrole-containing metabolites.

**Figure 6.**
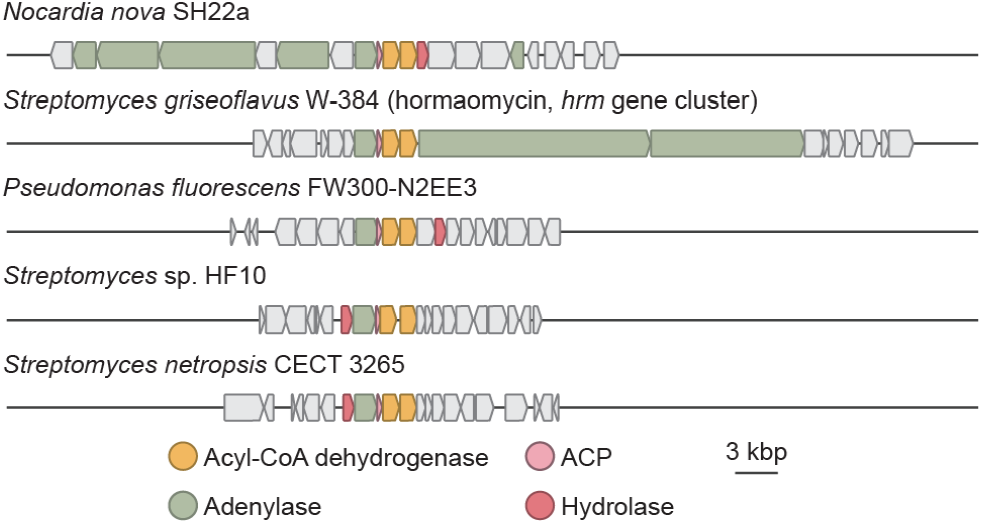
The *N*-hydroxypyrrole biosynthetic genes appear in many BGCs, suggesting this heterocycle is present in multiple undiscovered natural products.

### A heme-dependent guanidine *N*-oxygenase replaces the diiron HDO domain of SznF

We next sought to characterize chalkophomycin biosynthetic enzymes involved in diazeniumdiolate formation. Our stable isotope feeding experiments revealed L-arginine as the biosynthetic precursor of this functional group, suggesting a pathway for *N*-nitrosation that parallels that of streptozotocin. However, because the SznF homolog in chalkophomycin biosynthesis (ChmM) lacks the amino acid residues required for diiron binding in its HDO domain, we hypothesized that it would be unable to generate the L-dihydroxyguanidine intermediate required for *N*-nitrosation. Therefore, we proposed this pathway would use a different *N*-oxygenating enzyme. ChmN is annotated as a member of the “YqcI/YcgG uncharacterized protein family”. Recently, two members of this protein family, AglA and GntA, were shown to be heme-dependent guanidine *N*-oxygenases in argolaphos and guanitoxin biosynthesis, respectively (Figure 7A).^51,52^ Additionally, DcsA, a related enzyme required for D-cycloserine biosynthesis was found to bind heme, although its activity could not be confirmed *in vitro*.^53^ Although ChmN only shares 31% aa ID with AglA and 22% aa ID with GntA, we hypothesized that it could catalyze *N*-oxygenation of L-arginine, paralleling the transformation performed by SznF’s HDO domain. Notably, 107 genes encoding ChmN homologs colocalize with genes encoding SznF homologs predicted to have inactive HDO domains, further supporting this proposal (Figure S18).

**Figure 7.**
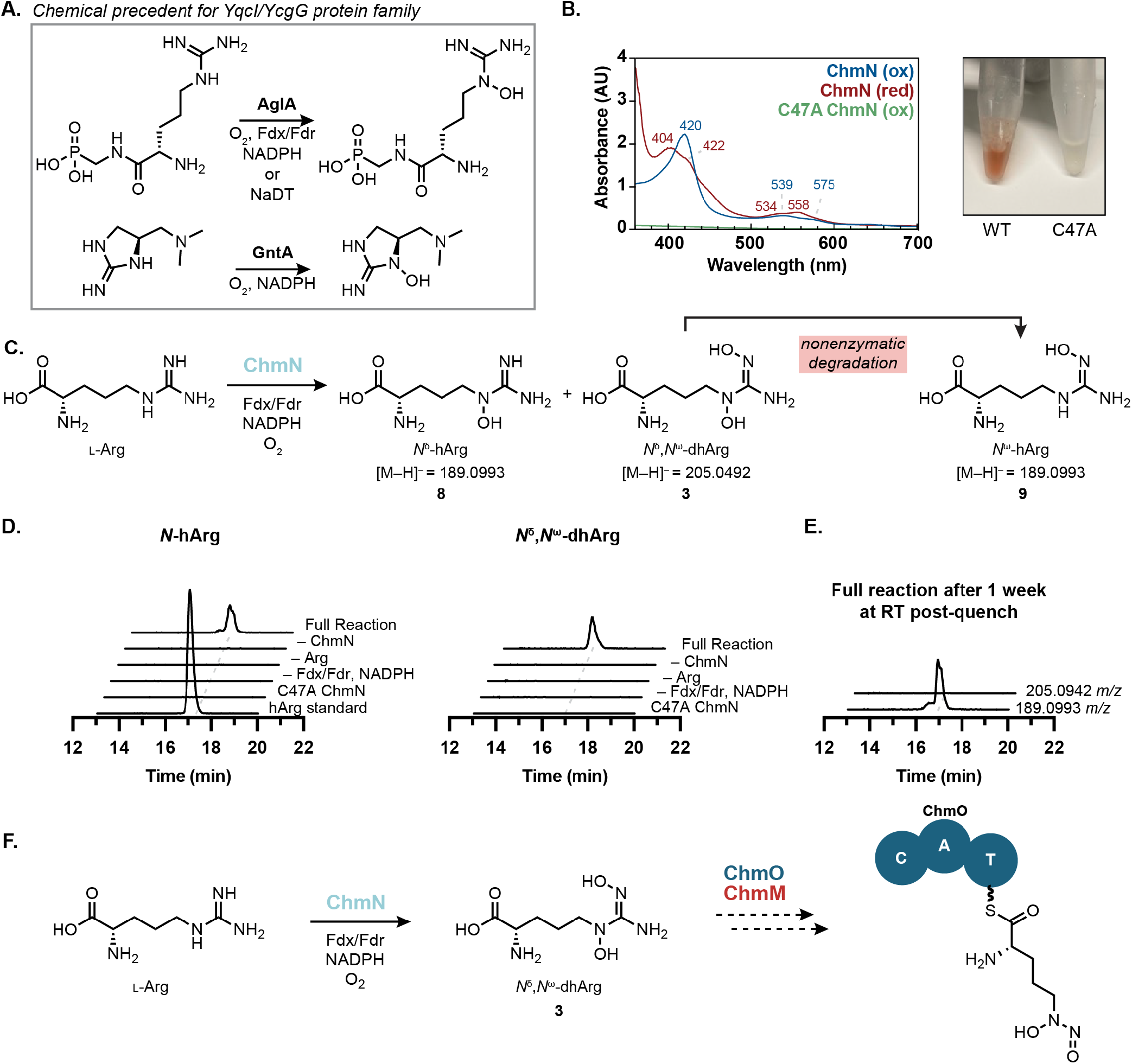
ChmN is a heme-dependent arginine *N*-oxygenase. (A) The “YqcI/YcgG uncharacterized protein family” has two biochemically characterized members, both of which catalyze guanidine *N*-oxygenation. (B) Purified ChmN has UV–vis spectroscopic features consistent with a thiolate-bound heme cofactor and is bright red. The C47A variant loses these features and all color, consistent with disruption of the heme-binding ligand. (C) ChmN catalyzes mono- and di-hydroxylation of L-arginine. (C) Incubation of ChmN with L-arginine and necessary redox cofactors results in the production of L-*N*^δ^-hydroxyarginine. The hArg standard is a mixture of *N*^δ^*-*hArg and *N*^ω^-hArg synthetic standards. (E) LC–MS samples for the full reaction were left at room temperature for one week and re-analyzed. Complete decomposition of *N*^δ^,*N*^ω^-dhArg was observed with a corresponding increase in concentration corresponding to a mixture of *N*^δ^-hArg and *N*^ω^-hArg. All EICs are graphed on the same y-axis range (0–2x10^5^ Ion Counts). (F) Proposed pathway for L-graminine biosynthesis involves the putative *N*-nitrosating enzyme ChmM and NRPS ChmO.

When recombinantly expressed in *E. coli*, ChmN bound heme, evidenced by characteristic UV–vis spectroscopic features and a bright red color (Figure 7B). These features and color were lost when the putative active site residue Cys47 was substituted with Ala, supporting its role as the axial heme-binding ligand. Moreover, when incubated with Larginine, NADPH, spinach ferredoxin, and ferredoxin reductase, ChmN catalyzes the production of *N*^δ^-hydroxyarginine (*N*^δ^-hArg, **8**) and *N*^δ^,*N*^ω^-dihydroxyarginine (*N*^δ^,*N*^ω^-dhArg, **3**) (Figure 7C, D). *N*^ω^hydroxyarginine (*N*^ω^-hArg, **9**) was also detected in reaction mixtures and was found to co-elute with *N*^δ^-hArg (**8**), evidenced by LC–MS/MS data and differential fragmentation patterns of the two hydroxyarginine isomers. We found that *N*^δ^-hArg (**8**) is produced more rapidly than *N*^ω^hArg (**9**) over a 3 h time course (Figure S19, Table S6). When supernatants of quenched reaction mixtures were left at room temperature for one week all *N*^δ^,*N*^ω^-dhArg (**3**) disappeared, while we saw an overall increase in the mass corresponding to both hydroxyarginine isomers (Figure 7E). Due to the correlation between *N*^δ^,*N*^ω^-dhArg (**3**) degradation and an increase in *N*^ω^-hArg concentration, we concluded that *N*^ω^hArg (**9**) is the degradation product of *N*^δ^,*N*^ω^-dhArg (**3**). Although purified recombinant ChmM has not yet displayed activity towards any of these arginine derivatives, these data support the proposal that heme-dependent enzymes have evolved to collaborate with SznF-like enzymes in the synthesis of *N*-nitrosated products (Figure 7F). This is the first experimental evidence linking the “YqcI/YcgG” enzymes that commonly appear in graminine-encoding gene clusters to the biosynthesis of this diazeniumdiolate-containing amino acid.

## DISCUSSION

Although *N*-hydroxypyrroles have been proposed to originate from amino acids such as L-leucine and L-proline, the enzymes used to construct this heterocycle have never been characterized. By genome mining for biosynthetic pathways involving *N*-nitrosation, we identified the chalkophomycin BGC and revealed enzymes involved in *N*-hydroxypyrrole and diazeniumdiolate assembly. While this manuscript was in revision, an identical chalkophomycin gene cluster was identified in the genome of *S*. sp. CB00271; however, no biochemical characterization of any enzymes was performed.^54^

We demonstrate *in vitro* that L-proline is adenylated by ChmJ and loaded onto the carrier protein ChmI before undergoing *N*-oxygenation by the flavin-dependent enzyme ChmG. Following *N*-oxygenation, ChmH catalyzes the formation of the final *N*-hydroxypyrrole via an FAD-dependent 2or 4-electron oxidation. This product is then hydrolyzed by ChmF to enable mobilization of the free *N*-hydroxypyrrole-2carboxylic acid (**2**) building block. Characterization of the adenylase ChmK reveals a route for incorporating additional hydroxylated aromatic building blocks into nonribosomal peptides. This knowledge will aid identification of BGCs that encode *N*-hydroxypyrrole-containing natural products, including novel metallophores.

Prior to this report, thiotemplated biosynthesis of functionalized pyrroles has followed a prescribed path of pyrrolidine to pyrrole oxidation prior to further elaboration. Halogen, methoxy, and methyl substituents have also been observed on the carbon atoms of pyrroles in natural products (Figure S20, S21). Biochemical characterization of the pentabromopseudilin biosynthetic pathway shows the strategy for thiotemplated halopyrrole biosynthesis from L-proline involving an initial flavin-dependent 4-electron oxidation of ACP-bound L-proline and followed by halogenation of the ACP-bound pyrrole.^39^ Methylpyrrole in clorobiocin is synthesized in a similar fashion, where carrier protein-bound L-proline is first oxidized to the pyrrole by ACAD CloN3.^55^ CloN6 has been proposed to perform a radical-based methylation of the pyrrole due to conservation of a cysteine-rich sequence motif found in the radical *S*-adenosylmethionine (SAM) protein superfamily, and genetic deletions of *cloN6* provide support for methylation occurring after pyrrole formation.^56,57^ C2-methylpyrrole and methyoxypyrrole have also been described to originate from distinct pathways.^58,49,59^

The characterization of enzymatic *N*-hydroxypyrrole formation represents a deviation from this established logic. Unlike the established order of proline oxidation followed by pyrrole functionalization, the Chm system catalyzes *N*-hydroxylation of proline prior to pyrrole formation. This order of events could perhaps reflect a strict requirement for the more nucleophilic nitrogen of the pyrrolidine to engage the presumed hydroperoxyflavin electrophile to achieve *N*-oxygenation.

This logic was not expected from prior analyses of the hormaomycin BGC.^37^ It was noted upon analysis of the *hrm* BGC that it did not reveal an obvious candidate for *N*-oxygenation, and in fact, it was incorrectly proposed that the ChmG and ChmH homologs, HrmN and HrmM, worked together to catalyze pyrrole formation prior to *N*-hydroxylation by an unknown enzyme. Not only does our work reveal the functions of these enzymes, but it corrects this mis-held assumption. Moreover, ChmG is, to the best of our knowledge, the first instance of both enzymatic proline *N*-oxygenation and *N*-oxygenation of a carrier protein-bound substrate by a flavin-dependent enzyme.

Using the newly elucidated genetic basis for *N*-hydroxypyrrole formation, we identified cryptic BGCs that construct this structural motif. Although *N*-hydroxypyrrole biosynthetic genes only co-occur with putative *N*-nitrosating enzymes in the *chm* BGC, these genes do appear near additional genes encoding siderophore transporters and NRPSs. In other BGCs, they are clustered with genes encoding non-assembly linetype biosynthetic enzymes. These BGCs are found in the genomes of many *Nocardia* and *Pseudomonas* that are opportunistic human pathogens. Given that competition for iron and other metals influences bacterial virulence, it is possible these putative metallophores may contribute to growth or pathogenicity.^60,61^ These BGCs present exciting opportunities for natural product discovery that may reveal new biological functions for *N*-hydroxypyrroles.

In addition to characterizing *N*-hydroxypyrrole biosynthesis, we discovered the activity of ChmN, a heme-dependent arginine *N*-oxygenase. Our results provide the first experimental insights into how L-arginine is transformed into L-graminine. Interestingly, we observed that genes encoding members of this family often co-occur in gene clusters alongside genes encoding SznF homologs predicted to lack a functional HDO domain. Therefore, we propose that ChmN and its relatives functionally replace the HDO domain’s role in *N*-nitrosation. This enzyme adds to our knowledge of the nascent YqcI/YcgG protein family, with heme-binding and guanidine *N*-oxygenation emerging as common features. Of note, this would be the first case in which a YqcI/YcgG enzyme catalyzes dihydroxylation and in which the hydroxyguanidine product motif is not observed in the final natural product structure, suggesting great potential for how these enzymes may be implemented in diverse biosynthetic pathways. This work substantially increases our understanding of this poorly characterized enzyme family which has >4,000 members (IPR014988).

Although the N–N bond-forming activity in chalkophomycin biosynthesis has yet to be reconstituted, ongoing efforts are directed towards this goal. The NRPS protein ChmO is the most likely candidate for incorporation of L-graminine into chalkophomycin. Interestingly, ChmO has a completely distinct A-domain substrate recognition sequence than that found in the NRPS enzymes predicted to activate and incorporate L-graminine in the gramibactin, megapolibactin, plantaribactin, gladiobactin, and tistrellabactin biosynthetic pathways (Table S2).^6, 17^ While this could suggest that multiple A-domain sequences are dedicated to L-graminine adenylation, it could also indicate a distinct pathway for L-graminine biosynthesis where the L-dihydroxyguinidine precursor is recognized by the NRPS and the diazeniumdiolate is formed on a protein-tethered substrate. *N*-nitrosation of a protein-based aminoacyl thioester substrate would be a dramatic expansion of activity of the SznF enzyme family. Moreover, the ability of these enzymes to generate both diazeniumdiolates and *N*-nitrosoureas raises questions about the mechanistic differences between ChmM and SznF. Specifically, enzymes that form L-graminine must promote carbon and nitrogen excision, while *N*-nitrosourea formation retains all atoms from the L-dihydroxyarginine substrate in the product.

Over the last two decades, genome mining has enabled the discovery of new microbial BGCs and natural products. This effort and previous studies have demonstrated the value of genome mining in identifying metallophore-encoding gene clusters by searching for putative *N*-nitrosating enzymes that construct a key metal-binding functional group. The discovery of the chalkophomycin biosynthetic pathway and the elucidation of *N*-hydroxypyrrole biosynthesis underscore how such efforts can also provide unanticipated opportunities for enzyme and metabolite discovery.

## EXPERIMENTAL SECTION

### Cultivation of *Streptomyces anulatus* ATCC 11523 for chalkophomycin production

*Streptomyces anulatus* ATCC 11523 was purchased from the American Type Culture Collection (ATCC). *S. anulatus* ATCC 11523 was grown on Mannitol Soy Agar (MS agar) (20 g/L D-mannitol, 20 g/L soybean flour, 20 g/L agar) for 9 days or until spores were produced. Spores were scraped using a sterile 10 µL inoculating loop and used to inoculate 30 mL of liquid TSB medium. Cultures were incubated at 30 °C with shaking at 220 rpm for 2 days. 5 mL of TSB starter culture was used to inoculate 50 mL of M2 Medium (15 g/L soluble starch, 5 g/L Pharmamedia, 100 mg/L CuSO_4_**•**5H_2_O, 5 mg/L NaI, 3 g/L CaCO_3_, pH 7) with 1.5 g (dry weight) activated HP20 resin (n = 6). Cultures were placed in a 30 °C incubator shaking at 220 rpm for 7 days.

After 7 days, the 50 mL *S. anulatus* ATCC 11523 cultures were decanted into conical tubes, and the cells and resin were pelleted by centrifugation (3,220 x g, 10 minutes). The supernatant was removed, and the pellet was washed 3x with Milli-Q water or until the supernatant was clear after centrifugation. 45 mL of MeOH was added to the pellet and left to incubate for 10 minutes, with gentle inversion every few minutes. The resin was pelleted again by centrifugation and the MeOH supernatant was concentrated using a rotary evaporator and further dried under vacuum overnight.

To the concentrated residue was added 1 mL of MeCN and 1 mL of H_2_O. 10 µL of resuspended material was diluted into 90 µL of 5% MeCN in H_2_O. The samples were spun at 16,100 x g for 10 minutes, and the supernatants were transferred into vials for analysis by LC–MS, using an Agilent Q-TOF 6530 equipped with an Dual AJS ESI source and a Kinetex C18 column (1.7 µm, 100 Å, 150 x 2.1 mm) flowing at a rate of 0.2 mL/min in a column compartment heated to 35 °C. Solution A was H_2_O + 0.1% formic acid, and Solution B was MeCN + 0.1% formic acid. The LC method was: 5% Solution B for 5 min; 5% to 95% Solution B over 25 min, 95% Solution B for 5 min, 95% to 5% over 1 min, hold at 5% for 10 min. The following parameters were used for the Q-TOF: Gas Temp 275 °C, Drying Gas 11 L/min, Nebulizer 35 psi, Sheath Gas Temp 275 °C, Sheath Gas Flow 11 L/min, VCap 3500 V, Nozzle Voltage 500 V.

Cultivation of *Streptomyces* sp. Root63, *Streptomyces* sp. Root1295, and *Lentzea flaviverrucosa* DSM 44664 for chalkophomycin production. *Streptomyces* sp. Root63, *Streptomyces* sp. Root1295, and *Lentzea flaviverrucosa* were purchased from the Leibniz Institute DSMZ. Each strain was grown on MS agar until sporulated, about 3 days. Spores were scraped from the plate using a 10 µL inoculating loop and used to inoculate 30 mL of liquid TSB medium. Cultures were grown for 3 days at 30 °C with shaking at 220 rpm until saturated. 750 µL of TSB starter culture was used to inoculate 25 mL of R2B medium supplemented with 100 mg/L Cu(II)SO_4_ • 5H_2_O (400 µM final concentration) (n = 5 per strain). Cultures were incubated at 30 °C with shaking at 220 rpm for one week, after which cells were pelleted (8,000 x g, 10 minutes) and the supernatants were filtered through a 0.2 µm filter. The filtrate was lyophilized to dryness. To the concentrated residue was added 500 µL of LC–MS grade MeCN and 500 µL of LC–MS grade water. The resuspended material was diluted 1:10 into water, and samples were spun at 16,100 x g for 10 minutes. The supernatants were transferred into vials for analysis by LC–MS. The same parameters for chalkophomycin detection from *S. anulatus* cultures were used.

### Heterologous expression of *chm* gene cluster in *Streptomyces coelicolor* M1152

A 5 mL culture of *E. coli* ET12567/pUZ8002 pDualP-chm was grown in LB with apramycin (50 µg/mL), kanamycin (50 µg/mL), and chloramphenicol (20 µg/mL) at 37 °C with shaking at 190 rpm for two days until saturated. Cells were passaged 1:100 in a 5 mL culture of LB with apramycin (50 µg/mL) and incubated at 37 °C with shaking until the OD_600_ reached 0.6, after which the cells were pelleted at 4000 x g for 5 minutes. The cells were washed twice with 5 mL LB without any antibiotics. After washing, the cells were resuspended in 500 µL of LB. To prepare the heterologous host, *Streptomyces coelicolor* M1152, for conjugation, 10 µL of spores from a frozen glycerol stock were added to 500 µL 2xYT medium. Spores were heat shocked at 50 °C for 10 minutes and allowed to cool before they were added to the 500 µL of concentrated *E. coli* cells. The mixture was briefly centrifuged at 8000 x g to pellet the cells. Supernatant was removed to leave ∼50 µL of liquid. The pelleted cells were resuspended by gently pipetting up and down, and they were plated on MS agar that contained 10 mM MgCl_2_ and incubated at 30 °C. 16 h after plating, the plate was overlayed with 0.5 mg nalidixic acid and 1.25 mg apramycin in 1 mL of water and then returned to incubate at 30 °C for 6 days. Successful exconjugants were re-streaked on MS agar with apramycin (50 µg/mL) and nalidixic acid (30 µg/mL). 20 mL of TSB medium was inoculated with spores from a plate of wild-type *S. coelicolor* M1152 or *S. coelicolor* pDualP-chm grown on MS agar with no antibiotic or apramycin (50 µg/mL) for the respective strains. TSB starter cultures of wild-type *S. coelicolor* contained no antibiotics and cultures of *S. coelicolor* pDualP-chm contained 50 µg/mL apramycin. After 3–4 days of shaking at 30 °C and 200 rpm, 750 µL of culture was used to inoculate 25 mL of R2B + 100 mg/L Cu(II)SO_4_ • 5H_2_O (n = 6). Cultures were left to incubate for 9 days at 30 °C with shaking 200 rpm, after which cells were pelleted at 8,000 x g for 10 minutes. Supernatants were passed through a 0.2 µm filter and lyophilized to dryness.

The concentrated culture supernatants were resuspended in 500 µL of LC–MS grade water and 500 µL of LC–MS grade MeCN and then diluted 1:10 in water. The samples were centrifuged at 16,100 x g for 10 minutes, and the sample supernatants were used for LC–MS analysis of chalkophomycin production. LC–MS parameters are identical to those listed above for detecting chalkophomycin in *S. anulatus* cultures.

### Stable isotope feeding in *Streptomyces* sp. Root63

*Streptomyces* sp. Root63 was struck out on MS agar from a frozen spore stock stored at –70 °C until sporulating. 20 mL of sterile TSB was inoculated with spores from this plate and incubated at 30 °C with shaking at 200 rpm for 3–4 days. 750 µL of the TSB seed culture was passaged into 25 mL of R2B + 100 mg/L Cu(II)SO_4_•5H_2_O and returned to the incubator at 30 °C with shaking at 200 rpm. After 48 h, 1 mM isotope-enriched substrate was added to cultures [^13^C_6_,^15^N_4_-l-arginine (99%) (n = 3), ^15^N_2_-l-ornithine (98%) (n = 3), ^15^N-sodium nitrite (98%) (n = 2), or ^15^N-proline (98%) (n = 3); Cambridge Isotope Laboratories]. After 5 d of growth, cultures were filtered through a 0.2 µm filter, and the filtrate was lyophilized to dryness. Lyophilized filtrate was resuspended in 500 µL of LC–MS grade MeCN and 500 µL of Milli-Q water. This solution was diluted 1:5 into Milli-Q water to prepare samples for LC–MS. Samples were analyzed by LC–MS on a ThermoFisher Orbitrap IQ-X equipped with an HESI source using a Kinetex C18 column (1.7 µm, 100 Å, 150 x 2.1 mm) flowing at a rate of 0.4 mL/min in a column compartment heated to 35 °C. Solution A was H_2_O + 0.1% formic acid, and Solution B was MeCN + 0.1% formic acid. The LC method was: 5% Solution B for 2 min; 5% to 80% Solution B over 27 min, 80% Solution B for 3 min, and re-equilibrated to 5% Solution B for 3 min. The following parameters were used for the Orbitrap detection: Resolution 60k, RF Lens 35%, Standard AGC Target, and Auto Maximum Injection Time Mode.

### General method for expression and purification of Chm enzymes

All proteins were expressed using BL21(DE3) *E. coli* or BAP1(DE3)^62^ *E. coli* and appropriate expression vectors (see Supporting Information for further details). An overnight culture of liquid LB medium supplemented with either 50 µg/mL kanamycin or 100 µg/mL ampicillin was inoculated from a frozen glycerol stock of *E. coli* harboring the expression vector for the desired protein and allowed to shake at 170 rpm and 37 °C. pET28a and pETDuet-1 were the two expression vectors used in this study. For each liter of protein expression culture (LB + antibiotic), 10 mL of the overnight culture was used for inoculation. The cultures were incubated at 37 °C with shaking at 180 rpm until they reached OD_600_ 0.4–0.6. Protein expression was induced by addition of IPTG (250 µM), and the cultures were returned to shaking with a lowered temperature at 16 °C for overnight expression.

The following day, cells were harvested by centrifugation at 6,730 x g. The cell pellet was resuspended in lysis buffer (50 mM HEPES, 500 mM NaCl, 10 mM MgCl_2_, pH 8) and sonicated. Lysate was clarified by centrifugation (18,000 x g, 35 min), and the clarified lysate was applied to a column of Ni-NTA resin (Qiagen and ThermoFisher) preequilibrated in wash buffer 1 (50 mM HEPES, 500 mM NaCl, 10 mM MgCl_2_, 20 mM imidazole, pH 8) for purification. The resin was washed with wash buffer 1, followed by wash buffer 2 (50 mM HEPES, 500 mM NaCl, 10 mM MgCl_2_, 60 mM imidazole, pH 8), before eluting the protein of interest with elution buffer (50 mM HEPES, 500 mM NaCl, 10 mM MgCl_2_, 200 mM imidazole, pH 8). The protein was buffer exchanged into storage buffer (20 mM HEPES, 50 mM NaCl, 10% w/v glycerol, pH 7.5) prior to flash freezing in liquid nitrogen. Protein aliquots were stored at –70 °C until needed.

## ASSOCIATED CONTENT

### Supporting Information

The Supporting Information is available free of charge on the ACS Publications website.

## AUTHOR INFORMATION

### Authors

**Anne Marie Crooke** Department of Chemistry and Chemical Biology, Harvard University, Cambridge, MA 02138, USA; https://orcid.org/0000-0002-1581-6104

**Anika K. Chand** Department of Chemistry and Chemical Biology, Harvard University, Cambridge, MA 02138, USA; https://orcid.org/0009-0006-3604-8550

**Zheng Cui** Department of Chemistry and Chemical Biology, Harvard University, Cambridge, MA 02138, USA; https://orcid.org/0000-0002-1503-7098

The authors declare no competing financial interests.

## Author Contributions

A.M.C and E.P.B. conceived of the study. A.M.C and E.P.B. wrote the manuscript with approval from all authors. A.K.C. assisted in characterization of ChmIJ activity. Z.C. expressed and purified ChmL. A.M.C. performed all remaining experiments and bioinformatic analyses.

## Funding Sources

This work was supported by an NSF Graduate Research Fellowship (DGE 2140743) to A.M.C., a postdoctoral fellowship from the National Institute of General Medical Sciences (1F32GM151795) to Z.C., and a grant from the National Institute of Health (5R01GM132564-04) to E.P.B. E.P.B. is an HHMI Investigator.

## ACKNOWLEDGMENT

The authors would like to thank Dr. Beverly Fu, Dr. Grace Kenney, and Miguel Aguilar Ramos for their insight and helpful discussions on experimental details. The authors also acknowledge Dr. Jared Mayers, Michelle Wang, and Katarina Pfeifer for providing feedback on the manuscript. We note that this Article is subject to HHMI’s Open Access to Publications policy. HHMI laboratory heads have previously granted a nonexclusive CC BY 4.0 license to the public and a sublicensable license to HHMI in their research articles. Pursuant to those licenses, the author-accepted manuscript of this Article can be made freely available under a CC BY 4.0 license immediately upon publication.

## Discovery of *N*-hydroxypyrrole-forming enzymes

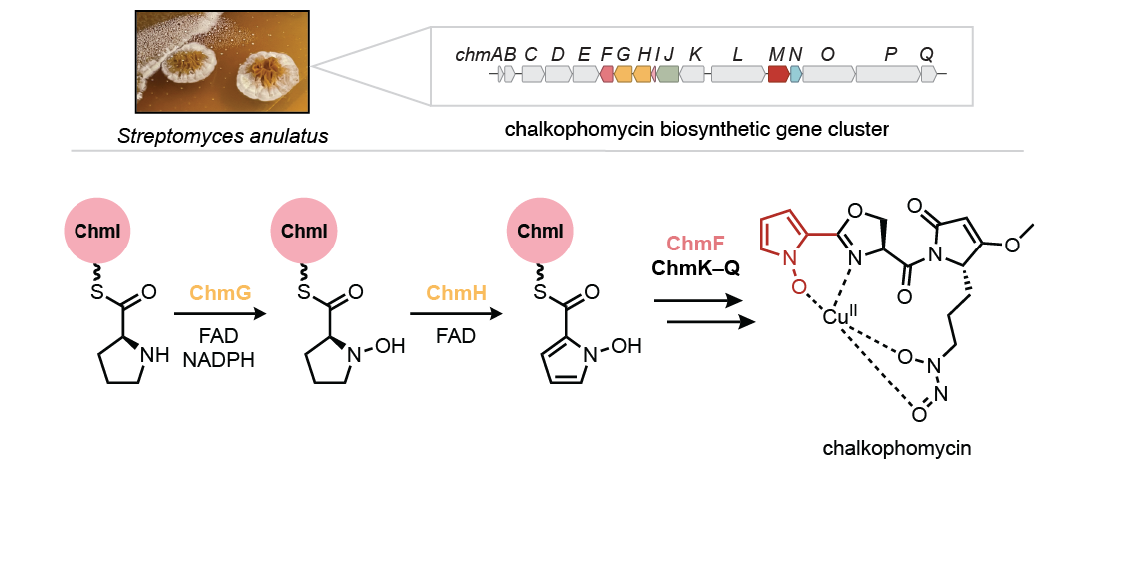

## Notes

### Competing Interest Statement

The authors have declared no competing interest.

### Summary of Updates

We include new experiments that 1) further support our findings demonstrating enzymatic formation of N-hydroxypyrrole in the chalkophomycin biosynthetic pathway and 2) demonstrate the role of two additional enzymes in this biosynthetic pathway for the mobilization of the N-hydroxypyrrole building block. We have also added more background to situate our findings in the context of the field.

