## Supporting Information for "Elucidation of chalkophomycin biosynthesis reveals *N*-hydroxypyrrole-forming enzymes"

Table of Contents


Materials and general methods ........ S1

General method for cloning of *chm* genes ........ S1

Codon optimized DNA sequences ........ S1

Expression and purification of proteins encoded by *chm* gene cluster ........ S2

N-His$_6$-ChmF ........ S2

N-His$_6$-ChmI ........ S3

N-His$_6$-ChmJ ........ S3

Co-expression of N-His$_6$-ChmI and ChmJ ........ S3

N-His$_6$-*Lfl*ChmG ........ S3

N-His$_6$-*Lfl*ChmH ........ S3

C-His$_6$-ChmN ........ S3

*N*-His$_6$-*Alfi*FIRed ........ S3

N-His$_6$-ChmL ........ S3

*In vitro* biochemical assays ........ S4

Loading of L-proline onto ChmI following adenylation by ChmJ ........ S4

*N*-hydroxypyrrole formation by *Lfl*ChmG and *Lfl*ChmH ........ S4

Release of *N*-hydroxypyrrole-2-carboxylic acid from ChmI by the hydrolase ChmF ........ S4

Guanidinyl *N*-oxygenation by ChmN ........ S5

Phylogenetic trees ........ S5

Synthesis of $N^{\delta}$-hydroxy-L-arginine ........ S5

Supplemental tables and figures ........ S8

Table S1: Putative function of proteins encoded by the *chm* biosynthetic gene cluster. ........ S8

Table S2: Substrate predictions for the adenylation domain-containing proteins ChmJ, ChmK, ChmL, and ChmO.$^{+}$ ........ S9

Table S4: Structural similarity search with ChmF using Dali Protein Structure Comparison Server (PDB25). ........ S11

Figure S1: SznF catalyzes guanidine *N*-oxygenation and *N*-nitrosation chemistry. ........ S12

Figure S2: Extracted ion chromatograms (EICs) for chalkophomycin production in native *chm* encoders. ........ S13

Figure S3: LC–MS/MS of 455.0497 *m/z* ion in *chm*-encoding bacteria. ........ S14

Figure S4: Identification of apo-chalkophomycin from *S*. sp. Root63 as a likely source fragment of $Cu^{II}$-bound chalkophomycin. ........ S15

Figure S5: LC–MS/MS of chalkophomycin ion for stable isotope feeding experiments ........ S16

Figure S6: Percent enrichment observed in stable isotope feeding experiments to determine chalkophomycin biosynthetic precursors. ........ S17

Table S5: Individual replicate and average values for percent enrichment observed in stable isotope feeding experiments. ....S17

Figure S7: The *N*-hydroxypyrrole functional group likely has several biosynthetic origins. ..S18

Figure S8. ChmJ adenylates proline for loading onto ChmI. ....S19

Figure S9: Multiple Sequence Alignment of ChmG and ChmH with a selection of known proline oxidases. ....S20

Figure S10: Phylogenetic tree provides insight into ChmG and ChmH function....S21

Figure S11: *Lfl*ChmH is inactive towards ChmI–proline....S22

Figure S12: $FADH_2$ availability influences *Lfl*ChmG product ratios....S23

Figure S13: Sequential addition of *Lfl*ChmH to ChmIJG reaction results in production of *N*-hydroxypyrrolyl-ChmI. ....S24

Figure S14: ChmF hydrolyzes the *N*-hydroxypyrrole product from ChmI. ....S25

Figure S15: ChmK adenylates N-hydroxypyrrole-2-carboxylic acid and transfers it onto the thiolation (T) domain of ChmL....S26

Figure S16: There is variety within *chmIJGH* homolog-containing gene clusters....S27

Figure S17: Structural alignment of ChmN, AglA, GntA, and DcsA AlphaFold predictions. .S28

Figure S18: Maximum likelihood phylogenetic tree shows conservation of cupin domain only SznF homologs with genes encoding for members of the YqcI/YcgG protein family. ....S29

Figure S19: LC–MS/MS analysis of ChmN time course. ....S30

Table S6: Ratios of 147:156 *m/z* fragment ions....S30

Figure S20: Selection of pyrrole-containing natural products discussed in this manuscript. S31

Figure S21: Strategies for functionalized pyrrole biosynthesis. ....S32

### Materials and general methods

Primers were supplied and gene sequencing was performed by Genewiz from Azenta Life Sciences. Codon-optimized genes were synthesized, and the corresponding expression vectors were assembled by Twist Biosciences. The heterologous expression construct (pDualP-chm) was prepared by Terra Bioforge (Varigen Biosciences). Q5 and Phusion Polymerases from New England Biosciences (NEB) were used for PCR amplification of target genes. Hifi DNA Assembly Master Mix from NEB was used for assembling protein expression vectors. The Zymoclean Gel DNA recovery kit was purchased from Zymo Research and the QIAquick PCR purification kit was purchased from Qiagen. BL21(DE3) *E. coli* (Invitrogen) and BAP1 *E.coli*[1] (lab stock) were used for protein expression. Protein concentrations were calculated using predicted extinction coefficients by the ExPASy ProtParam tool (https://web.expasy.org/protparam/) and measured absorbance at 280 nm. Absorbances were measured using either an Agilent 3500 or Thermo Fisher Scientific NanoDrop 2000 UV–vis spectrophotometer. Chemicals were purchased from Sigma-Aldrich and VWR. All extracted ion chromatograms (EICs) obtained on the Agilent Q-TOF 6530 were performed with a 10 ppm error range, and those obtained on the Orbitrap IQ-X were performed with a 5 ppm error range, unless otherwise stated.

### General method for cloning of *chm* genes

Genes were cloned from the pDualP-chm BAC.

Primers used for cloning (5' to 3')

| Gene | Forward primer | Reverse primer |
|---|---|---|
| *chmF* | TGGTGCCGCGCGGCAGCCATATGCCGGAGATCGAGCTG | TGGTGGTGGTGGTGCTCGAGTCAGGACGTCGGCGCC |
| *chmI* | CACCACAGCCAGGATCCGATGGAAGACGTCAATG | TTCTGTTCGACTTAAGCATCATGACACCGGGCTC |
| *chmJ** | GTATAAGAAGGAGATATACATATGAGCCGTCCCCC | GGCAGCAGCCTAGGTTAATCATTGACGTCTTCCATG |
| *chmJ*** | TGGTGCCGCGCGGCAGCCATATGAGCCGTCCCCC | TGGTGGTGGTGGTGCTCGAGTCATTGACGTCTTCCATG |
| *chmN* | CTTTAAGAAGGAGATATACCATGCGTATGTTGTCGGCG | GAGTGCGGCCGCAAGCTTGTCACCGGTGCCCTCACC |
| *chmL* | TGGTGCCGCGCGGCAGCCATATGGGAGACGTCGTGTTGG | TGGTGGTGGTGGTGCTCGAGTCAGGACTCGGCGAGC |

*primers used for untagged ChmJ co-expression with ChmI
**primers used for solo $His_6$-ChmJ expression

Primers used for site-directed mutagenesis for C47A ChmN

| Gene | Primer 1 | Primer 2 |
|---|---|---|
| *chmN* | CGCGCCGTTCCCC<u>GCC</u>ACCTTCGCGGTC | GACCGCGAAGGT<u>GGC</u>GGGGAACGGCGCG |

### Codon optimized DNA sequences

<u>*Lfl*ChmG</u>
ATGCGGACCATCGACGTGCACGATATGGGAGTAACGCTGAGACAAGTAGCTGAGGTAGCAGATGAGCTGC
GCCTGCTCGCGGCTGAAGTAGACGATGGGAGAGTCCCAGCGAGCCGCGGGTACGCGGCAGTTCGAAATGC
GGGGTTACTGCGGTTGCTGTTGCCACAATCCGCGGGTGGGTCGGGCTTATCCTACTTAGACTATACTAGA
GTGCTGGAAGAGCTCGCACTCGGGGACGCGAGTGCTGCCTTGGGATACAACATGCATAACGTGGCAATAG
GCGGGCTCTGCGAATCGGCAGAGCGCGACTTGCCACCCTCAGCTGCGGCTTTCAGACGGTGGGTGTTCGA
CGAGGTAGCTCATCGCGACCGCATGTTCGCTAGTGCTACCAGTGAAGTAGGCGGTGGCGCTAAACTCACA
GCGGTAAAAGCTACTTATCGGCGGACAGCTGACGGGTTCGTGCTGAATGGCCGCAAGTCGTTCGTTTCAT

TAGCTGGAATTGCCGATTACTACATAACCACAGCTCGGCGGGAAGACTCCGACGCAGCTGACGAGGTCTC
ACACTTCGTTGTTGCCAAGGACGACGCAGGGGTTAGTTTCGACGGCTTATGGGATGGTGCAGCCCTCGCA
GGAACCGCGACCTCGGCCATGACCCTTTCTAACGTGCACCTTGGTACCGATAGACTGTTCTTGGGAATAG
AAGGTATGAGCCTCTTCAAGCTTGTTCGGGAACCGCACTGGATGGTCTCTGGTTACATGGGTGCCTACCT
GGGTGTTGCCGAGTCTATTCTTCGCCACGTCACCACATTTGTTACCGCCGACGCAAAGCGCCGGGACAGT
GCAGTTGTGCGAGCTGAGGTAGGTCGCATGGCCGTTGACCTTGCAGCGACACGCGCCTTAGTTCATGCTG
CGGCTAGATTGGTGGATGAGAGAAAAGGGTCCGTTGAAGCCAACTCGGCCGTACACGCGGCCAAATACCG
CGTCGGAGAATCTACGACAACTCTGGCTCAAGCCGCAGTTCGTATTTGCGGGAGCACTGCCTTGCGACGA
TCTAATCCCCTGGAGCGTCTCCTGCGCGAGGCCTCATTTTGCAGCGTCATGCCCGCGAAACCAGATGAGT
GCTTAGAGTACGTTGGTAAGGCGAGACTTGGCTTCAATATGTTCGACTCTACGACTGTGGAATGGTGA

*Lfl*ChmH
ATGAACACGGGTGCGACTGATTTCACGGCTCGATGGGCAGGCATAGCGAGTTCTGCGTTGGCCGGTCTCA
GCACAGCTGACTTCCGAACACGGTGGAAGGCTTTAGCGGACTCAGGTGTTCTCGCTGGAGTTCGCGACGG
ACCGCGCGGGGTAGTGACCGACACCTTAGCCGCAATTGAAGGGCTGGGAATAGGCGGAACGTCTCCTGGT
CTTTGCTACGCGTTGACATCGCAGCATTTCGGGGTTCAGTCTCCTCTCCGTGCCGCCGGATCAGCAGCTT
GGGCGGCCTTAAGTGAAGGTGTGGAGCGAGGAGATACCTTGATGTGTCATGCCCTGACAGAAGAGGGCGG
CGGGTCGGACCCATTGTCAATGGCAACACGGGCGGAACCGGCAAACGGTGGCTGGGTCCTGACCGGTAGA
AAAGCCTTCGTGACTGCAGCTCCTATCGCGGACCACGCGTTGGTTTTCGCTCGAACGGATGTTGAACGAA
CTCCATTCGCCCTTAGCGCATTCTTAGTAGACCTCTCTAGCGCCGGCGTTAGCCGTTCAGAACCGTTTCC
AAAGAAGGCGTTGACCGATGTCCCGATGGGAGCCATCGATTTTGACTCTGTTCGTCTGGGTCCAGAGCAG
CTCGTGGGTCGTGAGGGTGCTGGGCTCGCCTTACTGACAACAACAACTTCTTGGGAGCGGGGCTTGCTGT
TGGGTTATGCACTCGGCCCTATGCGCCTCGTTTTAGACCGTACCGTTGAGTGGGCCCGCCGGCGAAAGCA
ATTCGGGCGGCCTATGGGTGCCTCTCACTTAGTCGCAGGTCGCCTGGCGGACATGGCGACACTGTTACAT
AGAAGTCGTACCTTAATTTTCGCCATGGCACAACGCTTCGACTCAGGAGAACCGGCTCGGCGTCTTGCGT
CCGAAGCCGCAATGACTAAGGTCTCCGTCTCAGAGGACCTCTTATCTTTCACACAGCATGCGGCAGCGCT
TGGCGGTGTACGAAGCTTTGTCAGTGATACCGGTTTAACCGCAGATCTGACGGACGCGATGGCCGCTGCG
GTCTACGCCGGTCCGAATGACCTTCTTCGTGTGAGTGTGGCGCGGGAGCTCGGCCTTCCGGTTGAGAATT
AA

*Aliivibrio fischeri* Flavin Reductase (*Alfi*FlRed)
ATGCCTATCAACTGTAAGGTGAAGAGCATCGAGCCCTTAGCTTGTAATACGTTTCGCATTTTATTACACC
CGGAGCAACCTGTTGCGTTTAAAGCAGGGCAATACCTGACGGTTGTAATGGGCGAAAAGGACAAGCGCCC
GTTTTCCATCGCGAGTTCTCCTTGTAGACATGAAGGGGAAATAGAGTTACATATAGGGGCAGCCGAACAC
AATGCTTATGCGGGTGAGGTTGTCGAATCCATGAAGTCTGCGCTTGAGACCGGTGGTGATATTTTGATAG
ACGCGCCTCACGGCGAGGCTTGGATCCGGGAAGACAGCGAGAGATCGATGTTATTGATAGCAGGTGGTAC
AGGCTTCTCGTACGTGAGAAGCATTCTGGACCACTGCATCAGTCAGAAGATCCAGAAACCCATATACTTA
TACTGGGGCGGTCGTGACGAGTGTCAGCTTTACGCTAAGGCGGAGCTTGAGACGATTGCCCAAGCGCATT
CCCACATTACATTTGTACCCGTAGTAGAGAAGAGCGAGGGTTGGACAGGCAAAACGGGTAACGTCTTAGA
AGCAGTAAAGGCTGACTTTAACTCCCTTGCAGACATGGACATTTATATCGCAGGGAGATTCGAGATGGCG
GGCGCCGCACGCGAGCAGTTCACGACTGAAAAGCAGGCCAAGAAAGAGCAACTGTTCGGAGACGCCTTCG
CTTTCATTTGA

### Expression and purification of proteins encoded by *chm* gene cluster

The general method used for protein expression and purification is detailed in the main text Experimental Section. Below are deviations from the general method for each protein.

N-His$_6$-ChmF
pET28a was used for expression of N-His$_6$-ChmF, and kanamycin (50 µg/mL) was used for antibiotic selection. Expression was induced at $OD_{600}$ 0.4–0.6.

N-$His_6$-ChmI
pETDuet-1 was used for expression of N-$His_6$-ChmI, and ampicillin or carbenicillin (100 µg/mL) was used for antibiotic selection. Protein expression was induced at $OD_{600}$ 0.1–0.24.

N-$His_6$-ChmJ
pET28a was used for expression of N-$His_6$-ChmJ, and kanamycin (50 µg/mL) was used for antibiotic selection. Expression of ChmJ was induced at $OD_{600}$ 0.3.

Co-expression of N-$His_6$-ChmI and ChmJ
pETDuet-1 was used for expression of N-$His_6$-ChmI and untagged ChmJ in BAP1 *E. coli*(DE3). Ampicillin (100 µg/mL) was used as the appropriate antibiotic. Protein expression was induced at $OD_{600}$ 0.4–0.9. N-$His_6$-ChmI was able to pulldown untagged ChmJ by Ni-affinity chromatography. The purified protein was buffer exchanged using a PD-10 desalting column with standard storage buffer.

N-$His_6$-*Lfl*ChmG
pET28a was used for expression of N-$His_6$-*Lfl*ChmG, and kanamycin (50 µg/mL) was used for antibiotic selection. 100 µM riboflavin was added to protein expression cultures at the time of IPTG induction and the lysis buffer contained 1 mM FAD. The wash and elution buffers had an increased salt concentration of 300 mM NaCl and contained 10% w/v glycerol, and the purified protein was buffer exchanged using a spin concentrator into storage buffer (20 mM HEPES, 300 mM NaCl, 10% glycerol, pH 8).

N-$His_6$-*Lfl*ChmH
pET28a was used for expression of N-$His_6$-*Lfl*ChmH, and kanamycin (50 µg/mL) was used for antibiotic selection.100 µM riboflavin was added to protein expression cultures at the time of IPTG induction and the lysis buffer contained 1 mM FAD. The wash and elution buffers had an increased salt concentration of 300 mM NaCl and contained 10% w/v glycerol, and the purified protein was buffer exchanged using a spin concentrator into storage buffer (20 mM HEPES, 300 mM NaCl, 10% glycerol, pH 8).

C-$His_6$-ChmN
pET28a was used for expression of C-$His_6$-ChmN, and kanamycin (50 µg/mL) was used for antibiotic selection. In addition to the standard method for protein expression, 1 mM 5-aminolevulinic acid and 1 mM ferrous ammonium sulfate were added to the expression cultures at the time of inoculation. Elution buffer contained 500 mM imidazole.

*N*-$His_6$-*Alfi*FlRed
pET28a was used for expression of N-$His_6$-*Alfi*FlRed, and kanamycin (50 µg/mL) was used for antibiotic selection. Expression of *Alfi*FlRed was induced at $OD_{600}$ 0.8. 100 µM riboflavin was added to protein expression cultures at the time of IPTG induction and the lysis buffer contained 1 mM FAD. The same wash, elution, and storage buffers used for ChmG and ChmH were used.

N-$His_6$-ChmL
pET28a was used for expression of N-$His_6$-ChmL from BAP1(DE3) *E. coli*, and kanamycin (50 µg/mL) was used for antibiotic selection. For each liter of protein expression culture (Terrific Broth with 35 µg/mL kanamycin), 10 mL of the overnight culture (prepared using the general method described in the main text) was used for inoculation. The cultures were incubated at 12 °C with

shaking at 200 rpm until they reached $OD_{600}$ 0.4–0.6. Protein expression was induced by the addition of IPTG (150 µM), and the cultures were returned to shaking at 12 °C for 48 h expression.

#### ***In vitro*** **biochemical assays**

All reagent stock solutions were prepared in the enzyme assay reaction buffer, the composition of which is detailed in each section. Enzyme stock solutions were made up of the enzyme storage buffer, which is outlined in the main text Experimental Section or Supporting Information (*Lfl*ChmG, *Lfl*ChmH). All *in vitro* reactions were carried out in at least three biological replicates.

Loading of ʟ-proline onto ChmI following adenylation by ChmJ

In a 50 µL reaction mixture, co-purified ChmI–ChmJ (20 µM) was combined in reaction buffer (50 mM HEPES, 200 mM NaCl, 10 mM $MgCl_2$, pH 8) with ATP (5 mM) and ʟ-proline (1 mM). The reaction mixtures were incubated at room temperature for 1 h. Reaction mixtures were frozen and stored at –80 °C until analysis.

An Agilent Q-TOF 6530 equipped with a Dual AJS ESI source was used for LC–MS analysis. A Dikma BioBond C4 column (5 µm, 5 x 4.6 mm) flowing at a rate of 0.5 mL/min was used. Solution A was $H_2O$ + 0.1% formic acid, and Solution B was MeCN + 0.1% formic acid. The LC method was: 5% Solution B for 3 min; 5% to 95% Solution B over 25 min, 95% Solution B for 2 min, 95% to 5% over 3 min, and 5% for 3 min. The following parameters were used for the Q-TOF: Gas Temp 325 °C, Drying Gas 10 L/min, Nebulizer 35 psi, Sheath Gas Temp 275 °C, Sheath Gas Flow 11 L/min, VCap 4000 V, Nozzle Voltage 1000 V.

*N*-hydroxypyrrole formation by *Lfl*ChmG and *Lfl*ChmH

To prepare ChmI-ʟ-proline, *apo* N-$His_6$-ChmI (20 µM), N-$His_6$-ChmJ (10 µM), Sfp (5 µM), Coenzyme A (1 mM), ATP (5 mM), and ʟ-proline (250 µM) were mixed in 8 µL of reaction buffer (50 mM HEPES, 200 mM NaCl, 10 mM $MgCl_2$, pH 8) and incubated for 1.5 h. Following the initial incubation, *Lfl*ChmG (20 µM), FAD (100 µM), NADPH (1 mM), and *Lfl*ChmH (20 µM) were added to the reaction mixture. Additional buffer was added to bring the final reaction volume to 50 µL.

After a 30 minute incubation, samples were diluted with 100 µL of ice-cold water and centrifuged at 16,100 x g for 10 minutes or passed through a 0.2 µm filter. Supernatants were removed or filtrate was collected for analysis by LC–MS.

The same mass spectrometry parameters were used as in the ChmI and ChmJ activity assay described above.

Note: for samples including *Alfi*FIRed (20 µM), the assay was performed in the same manner, and this component was added at the same time as *Lfl*ChmG. For controls with superoxide dismutase (100 U/mL) and catalase (35 U/mL), these enzymes were added to the reaction at the same time as *Lfl*ChmG. Both superoxide dismutase and catalase were purchased from Sigma Aldrich.

Release of *N*-hydroxypyrrole-2-carboxylic acid from ChmI by the hydrolase ChmF

Reaction mixtures were prepared in an identical manner to *Lfl*ChmG and *Lfl*ChmH assays with the addition of ChmF (10 µM) to all the other components for *N*-hydroxypyrrole formation. Samples for LC–MS analysis of the ChmI product was prepared in an identical manner as above and were analyzed using the same method on the Agilent Q-TOF 6530. For analysis of hydrolyzed products, the reaction was quenched with equal reaction volume of LC–MS grade MeCN and cooled on ice for 10 minutes. The samples were then centrifuged for 10 minutes at 16,100 x g, and supernatants were removed for analysis.

The ChmI protein product was analyzed by LC–MS using the same parameters as above. For analysis of the hydrolyzed product, a Kinetex C18 column (1.7 µm, 100 Å, 150 x 2.1 mm) flowing at a rate of 0.2 mL/min in a column compartment heated to 35 °C was used. Solution A was $H_2O$ + 0.1% formic acid, and Solution B was MeCN + 0.1% formic acid. The LC method was: 5% Solution B for 5 min; 5% to 95% Solution B over 15 min, 95% Solution B for 5 min, 95% to 5% over 1 min, hold at 5% for 10 min. The following parameters were used for the Q-TOF: Gas Temp 275 °C, Drying Gas 11 L/min, Nebulizer 35 psi, Sheath Gas Temp 275 °C, Sheath Gas Flow 11 L/min, VCap 3500 V, Nozzle Voltage 500 V.

Guanidinyl *N*-oxygenation by ChmN

For a 100 µL reaction in buffer (50 mM HEPES, 150 mM NaCl, pH 7.4), L-arginine (500 µM), spinach ferredoxin (5 µg), and spinach ferredoxin reductase (0.02 U), and NADPH (1 mM) were combined prior to addition of ChmN (10 µM). Reaction mixtures were incubated for 3 h at room temperature. 100 µL of MeCN was added to precipitate ChmN, and the reaction mixtures were placed on ice for 10 minutes, followed by centrifugation at 16,100 x g for 10 minutes at 4 °C. Supernatants were removed for analysis by LC–MS on an Agilent Q-TOF 6530.

A Cogent Diamond Hydride column (4 µm, 100 Å, 150 x 10 mm) was used for analyte separation at a flow rate of 0.5 mL/min. Solution A was $H_2O$ + 0.1% formic acid, and Solution B was MeCN + 0.1% formic acid. The LC method was: 90% Solution B for 1 min; 90% to 30% Solution B over 19 min, 30% Solution B for 1 min, 30% to 90% over 4 min, hold at 90% for 10 min. The following parameters were used for the Q-TOF: Gas Temp 300 °C, Drying Gas 11 L/min, Nebulizer 35 psi, Sheath Gas Temp 275 °C, Sheath Gas Flow 11 L/min, VCap 3500 V, Nozzle Voltage 500 V.

### Phylogenetic trees

Sequences for maximum likelihood phylogenetic trees were first aligned using MUSCLE in Geneious. IQ-TREE[2] was used for construction of the phylogenetic tree. Ultrafast bootstrap[3] was used with 1000 replicates. Trees were visualized using Interactive Tree of Life (iTOL)[4]. The tree of SznF-like enzymes contains the same protein sequences as the SSN in Main Text Figure 1. The tree of Acyl-CoA dehydrogenases was retrieved from the reviewed members of the InterPro family IPR006091.

### Synthesis of $N^{\delta}$-hydroxy-L-arginine

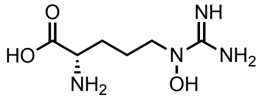


$N^{\delta}$-hydroxy-L-arginine was synthesized as previously described but with use of a Cbz protecting group instead of Boc.[5] $N^{\alpha}$-Cbz-$N^{\delta}$-hydroxy-L-ornithine HCl (69 mg, 216 µmol, 1 equiv.) was dissolved in anhydrous DMF (270 µL) under argon in an oven-dried flask. $Et_3N$ (30 µL, 499 µmol, 2.3 equiv.) was added in a dropwise fashion followed by 1*H*-pyrazole-1-carboxamidine hydrochloride (44 mg, 303 µmol, 1.4 equiv.) portion-wise. The reaction mixture was left to stir for 40 h at room temperature. The reaction was stopped by concentration, and the crude mixture was purified by automated flash chromatography using a Biotage Selekt with a 12 g C18 column. A two-step gradient of 2–10% acetonitrile in water, followed by 10–60%, and an isocratic hold at 60% acetonitrile in water was used to afford 30 mg *N*$^{\alpha}$-Cbz-*N*$^{\delta}$-hydroxy-L-arginine.

For removal of the Cbz protecting group, *N*$^{\alpha}$-Cbz-*N*$^{\delta}$-hydroxy-L-arginine (30 mg, 94 µmol) was dissolved in MeOH (3 mL) and a small spatula tip of both Pd/C and PdOH/C was added. The atmosphere was evacuated and replaced with $H_2$ using a balloon. The reaction mixture was left to stir at room temperature overnight. The following day, the reaction mixture was filtered over celite pre-wet with MeOH. The celite was rinsed thoroughly with MeOH and water to recover 19 mg of product that was used without further purification, as roughly a 1:1 a mixture of *N*$^{\delta}$-hydroxy-L-arginine and L-arginine. Spectral data has been reported for this compound in $CH_3OD$, and here we record the data for the mixture of *N*$^{\delta}$-hydroxy-L-arginine and L-arginine in $D_2O$. $^{1}$H NMR ($D_2O$, 400 MHz) δ 1.67–1.94 (m, 4H), 3.42 (dt, *J* = 136, 4 Hz, 1H), 3.59 (t, *J* = 4 Hz, 1H), 3.76 (q, *J* = 5 Hz, 1H). $^{13}$C NMR ($D_2O$, 101 MHz) δ 21.8, 27.4, 50.1, 54.3, 154.4, 174.7. Note: Peaks on the spectra below and not listed above are believed to arise from trace contaminants and MeOH.

**NMR spectra for *N*$^{\delta}$-hydroxy-L-arginine and L-arginine mixture. A.** $^{1}$H spectrum (recorded in $D_2O$ at 400 MHz). **B.** $^{13}$C spectrum (recorded in $D_2O$ at 101 MHz).

**A.**

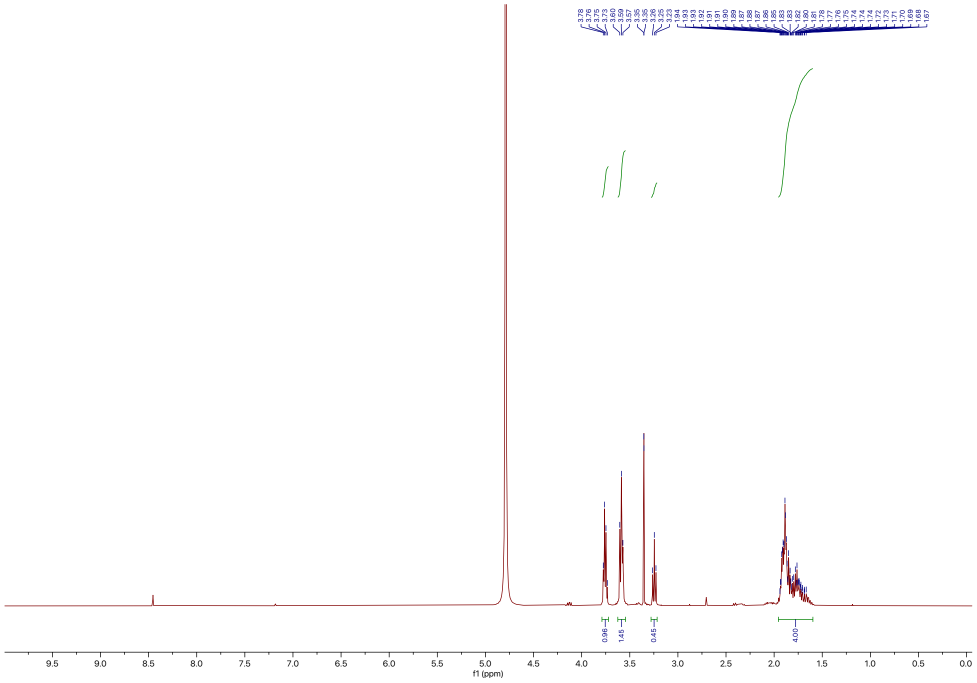

**B.**

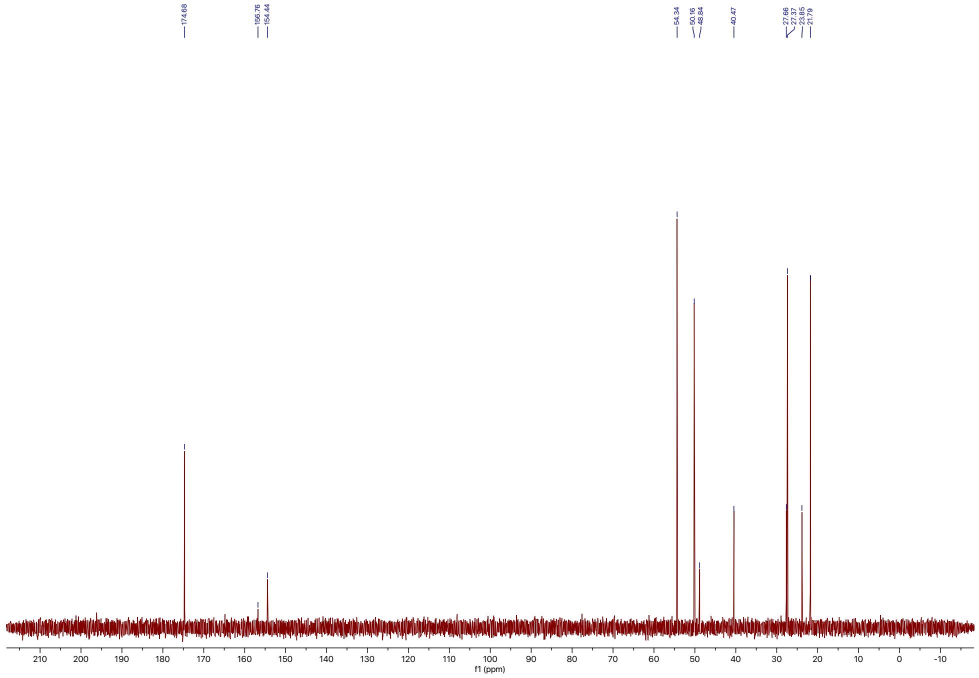

174.68
156.76
154.44
54.34
50.16
48.84
40.47
27.66
27.37
23.85
21.79
f1 (ppm)

### Supplemental tables and figures

**Table S1: Putative function of proteins encoded by the *chm* biosynthetic gene cluster.** Closest characterized homologs were identified through BLAST searches in the UniProtKB-SwissProt database.[+]

| Protein | Size (aa) | Putative function | Closest characterized homolog (% AA ID) | UniProtKB |
|---|---|---|---|---|
| ChmA | 116 | LuxR transcriptional regulator | Oxygen regulatory protein NreC (44%) | Q7A029 |
| ChmB | 242 | TetR/AcrR transcriptional regulator | Tetracycline repressor protein class C (28%) | P03039 |
| ChmC | 478 | MFS Transporter | Puromycin resistance protein pur8 (47%) | P42670 |
| ChmD | 595 | ABC Transporter | Mycobactin import ATP-binding/permease protein IrtA (43%) | G7CBF5 |
| ChmE | 582 | ABC Transporter | Mycobactin import ATP-binding/permease protein IrtB (38%) | G7CBF6 |
| ChmF | 264 | α/β hydrolase | Protein ABHD11 (26%), Proline iminopeptidase (31%) | Q8NFV4, P46542 |
| ChmG | 395 | ACAD: *N*-oxygenation | Putative acyl-CoA dehydrogenase YdbM (27%) | P96608 |
| ChmH | 383 | ACAD: pyrrole formation | L-prolyl-[peptidyl-carrier protein] dehydrogenase (32%) | Q4KCY6 |
| ChmI | 92 | Carrier protein | Lincomycin biosynthesis protein LmbN (carrier protein) (30%) | Q54367 |
| ChmJ | 517 | Adenylate-forming enzyme (Pro) | L-proline-[L-prolyl-carrier protein] ligase (33%) | Q4KCY5 |
| ChmK | 543 | Adenylate-forming enzyme (*N*-hydroxylpyrrole-2-carboxylic acid) | 2,3-dihydroxybenzoate-AMP ligase (52%) | P40871 |
| ChmL | 1180 | NRPS T-Cy-A-T (Ser) | Phenyloxazoline synthase MbtB (49%) | Q1B6A7 |
| ChmM | 469 | *N*-nitrosating | Acireductone dioxygenase (24%) | B1XPT2 |
| ChmN | 263 | Heme-dependent arginine *N*-oxygenase | Uncharacterized protein YqcI/YcgG (42%) | P45944 |
| ChmO | 1152 | NRPS C-A-T (?) | Phosphinothricin tripeptide synthase PhsC (36%) | D9XF47 |
| ChmP | 1414 | PKS KS-AT-?-T (Mal-CoA) | Phenolphthiocerol/phthiocerol polyketide synthase subunit E (43%) | P9WQE1 |
| ChmQ | 339 | *O*-methyltransferase | Mitomycin biosynthesis 6-*O*-methyltransferase (37%) | Q9X5T6 |

[+] analysis performed Summer 2022 and results may have changed since

**Table S2: Substrate predictions for the adenylation domain-containing proteins ChmJ, ChmK, ChmL, and ChmO.+**

| Protein | A-domain residues | antiSMASH (aS) | PRISM | Univ. of Maryland NRPS Predictor (UM) | Actual Substrate (biochemical validation or structural inference) |
|---|---|---|---|---|---|
| ChmJ | D-L-F-Y-A-A-I-V-C-K (aS)<br>D-L-F-Y-A-A-L-V (UM) | – | L-proline | – | L-proline |
| ChmK | T-M-P-A-Q-G-V-L-C-K (aS) | Dihydroxybenzoate | Salicylic acid | – | *N*-hydroxypyrrole-2-carboxylic acid |
| ChmL | D-L-Y-N-I-G-L-I-H-K (aS)<br>D-L-Y-N-I-G-L-I (UM) | L-cysteine | L-cysteine | L-threonine | L-serine |
| ChmO | D-I-N-Y-W-G-G-I-G-K (aS)<br>D-I-N-Y-W-G-G-I (UM) | L-ornithine | L-serine | L-*N*-hydroxyformyl ornithine | N.d. |

* "–" denotes that a domain or specific substrate-predicting residues were not recognized by the corresponding database
+ Substrate prediction was carried out in Fall 2021 and results may have changed since

Table S3: Structural similarity search with ChmG using Dali Protein Structure Comparison Server (PDB25).

| Result | PDB Code-Chain | Z-score | rmsd | lali | nres | %id | PDB Description |
|---|---|---|---|---|---|---|---|
| 1 | 4kcf-A | 41.5 | 2.4 | 366 | 407 | 24 | FAD-DEPENDENT OXIDOREDUCTASE |
| 2 | 7s7g-A | 36.5 | 2.7 | 358 | 571 | 19 | VERY LONG-CHAIN SPECIFIC ACYL-COA DEHYDROGENASE |
| 3 | 3d9d-A | 35 | 2.9 | 364 | 432 | 16 | NITROALKANE OXIDASE |
| 4 | 2or0-B | 34.8 | 3 | 358 | 409 | 22 | HYDROXYLASE |
| 5 | 4xvx-B | 33.7 | 2.7 | 340 | 373 | 17 | ACYL-[ACYL-CARRIER-PROTEIN] DEHYDROGENASE MBTN |
| 6 | 4rm7-A | 32.2 | 3.4 | 337 | 375 | 20 | ACYL-COA DEHYDROGENASE |
| 7 | 8hk0-B | 31.9 | 3 | 345 | 379 | 22 | DEHYDROGENASE |
| 8 | 6es9-A | 30.5 | 2.3 | 359 | 545 | 19 | ACYL-COA DEHYDROGENASE |
| 9 | 3owa-C | 28.6 | 2.3 | 359 | 587 | 19 | ACYL-COA DEHYDROGENASE |
| 10 | 8hk0-C | 25.7 | 4.2 | 314 | 346 | 21 | DEHYDROGENASE |

**Table S4: Structural similarity search with ChmF using Dali Protein Structure Comparison Server (PDB25).**

| Result | PDB Code-Chain | Z-score | rmsd | lali | nres | %id | PDB Description |
|---|---|---|---|---|---|---|---|
| 1 | 4opm-A | 26.9 | 2.2 | 237 | 299 | 19 | LIPASE |
| 2 | 1u2e-A | 26.5 | 2.4 | 239 | 286 | 18 | 2-HYDROXY-6-KETONONA-2,4-DIENEDIOIC ACID |
| 3 | 3e3a-B | 26.1 | 2.8 | 225 | 275 | 21 | POSSIBLE PEROXIDASE BPOC |
| 4 | 4q3l-B | 25.8 | 3 | 238 | 280 | 19 | MGS-M2 |
| 5 | 3nwo-A | 25.8 | 2.5 | 229 | 299 | 24 | PROLINE IMINOPEPTIDASE |
| 6 | 4pw0-A | 25.5 | 2.4 | 231 | 275 | 17 | ALPHA/BETA HYDROLASE FOLD PROTEIN |
| 7 | 5a62-A | 25.4 | 2.7 | 237 | 272 | 20 | PUTATIVE ALPHA/BETA HYDROLASE FOLD PROTEIN |
| 8 | 4qes-A | 24.8 | 3.1 | 244 | 441 | 21 | NON-HAEM BROMOPEROXIDASE BPO-A2, MATRIX PROTEIN 1 |
| 9 | 4g8b-A | 24.4 | 2.8 | 237 | 277 | 19 | ALPHA/BETA HYDROLASE FOLD PROTEIN |
| 10 | 4x00-A | 24.1 | 2.7 | 235 | 273 | 20 | PUTATIVE HYDROLASE |

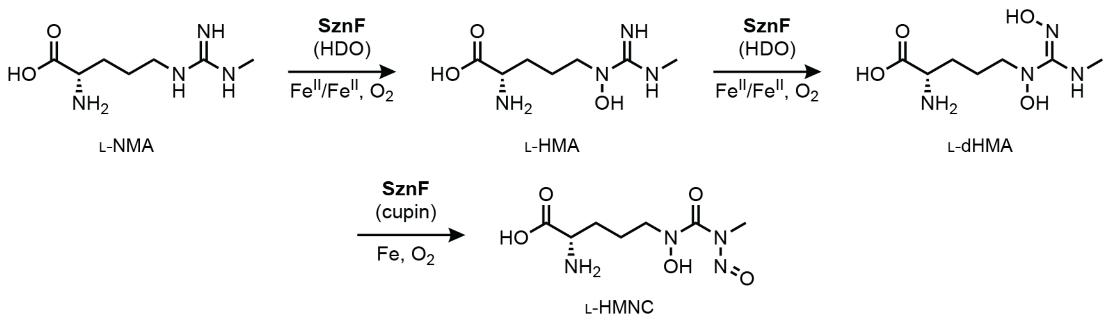


**Figure S1: SznF catalyzes guanidine *N*-oxygenation and *N*-nitrosation chemistry.** L-$N^{\omega}$-methylarginine (L-NMA); L-$N^{\delta}$-hydroxymethylarginine (L-HMA); L-$N^{\delta}$-hydroxy-$N^{\omega}$-hydroxy-$N^{\omega}$-methylarginine (L-dHMA); L-$N^{\delta}$-hydroxy-$N^{\omega}$-methyl-$N^{\omega}$-nitrosocitrulline (L-HMNC).

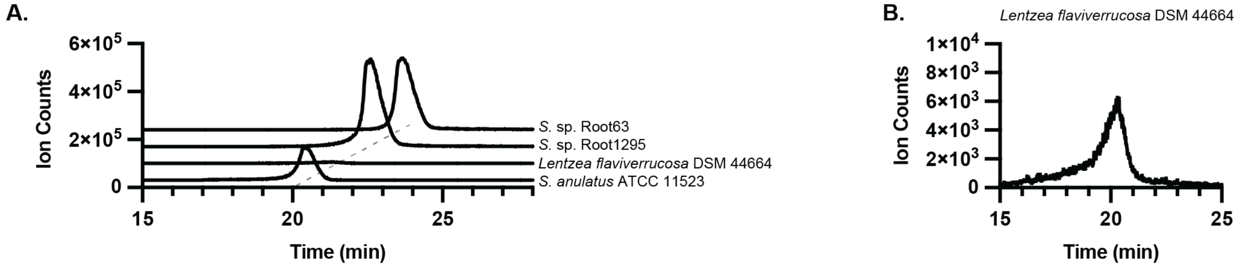


**Figure S2: Extracted ion chromatograms (EICs) for chalkophomycin production in native *chm* encoders. A.** EIC for chalkophomycin $[M+Cu^{II}–H]^+$ = 455.0497 (10 ppm error) for cultures of *S. anulatus* ATCC 11523 grown in M2 medium and *L. flaviverrucosa*, *S.* sp. Root1295, and *S.* sp. Root63 grown in R2B medium + 100 mg/L Cu(II)$SO_4$ • $5H_2O$. **B.** EIC of *L. flaviverrucosa* from panel A shown on smaller range y-axis.

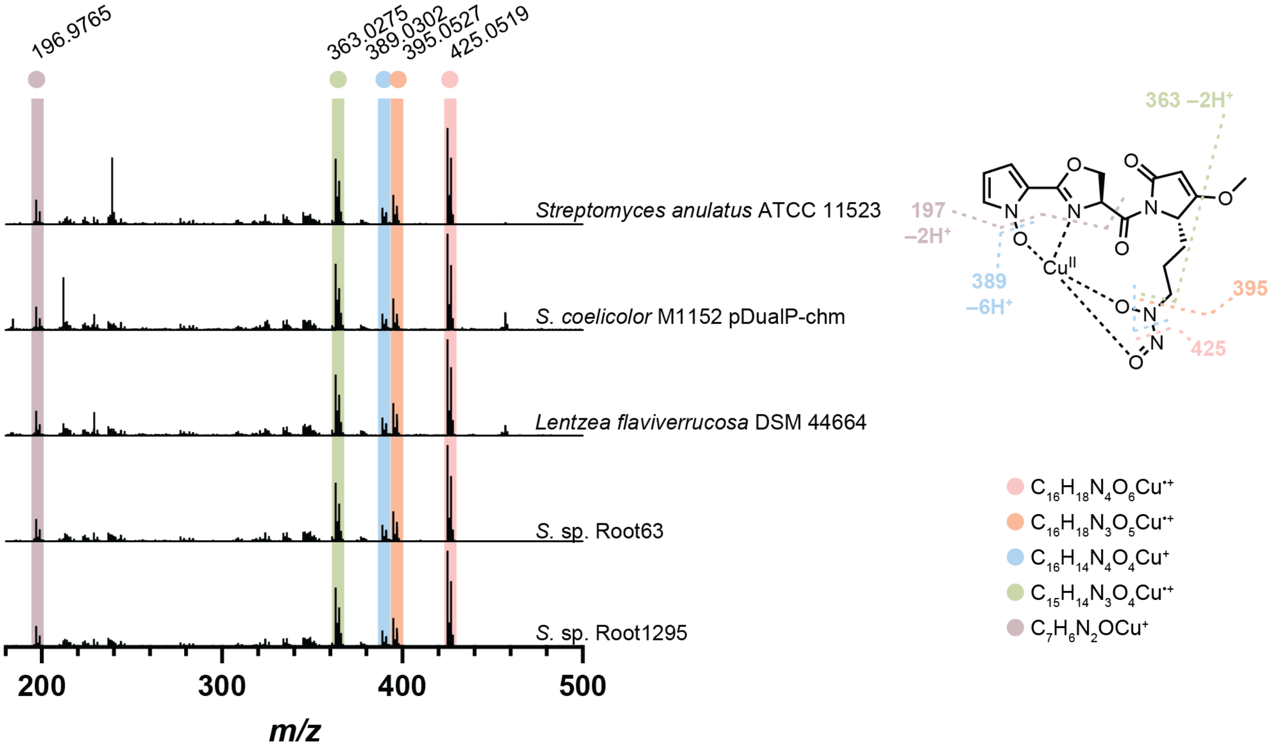


**Figure S3: LC–MS/MS of 455.0497 *m/z* ion in *chm*-encoding bacteria.** MS/MS fragmentation of 455.0497 *m/z* ion from cultures of *chm*-encoding bacteria. Diagnostic fragments are highlighted and shown with the most probable molecular formula. The most probable fragmentation mode for each molecular formula is depicted on the chalkophomycin structure. Data was acquired on an Agilent Q-TOF with the following MS/MS parameters: Frag=100.0V CID@20.

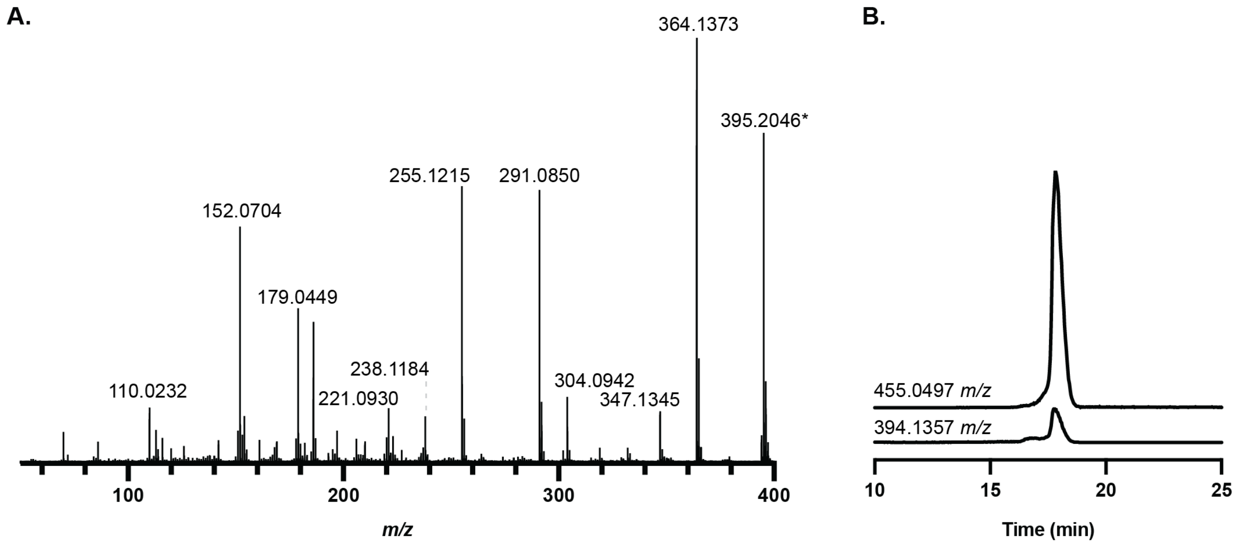


**Figure S4: Identification of apo-chalkophomycin from *S*. sp. Root63 as a likely source fragment of $Cu^{II}$-bound chalkophomycin.** A. LC–MS/MS of 394.1357 *m/z* ion, which matches previously reported chalkophomycin MS/MS data.[6] B. Retention times of EICs for $Cu^{II}$-chalkophomycin (455.0497 *m/z*) and apo-chalkophomycin (394.1357 *m/z*) are identical, suggesting the apo-chalkophomycin mass feature is a source fragment of the $Cu^{II}$-chalkophomycin metabolite. *Note that the 395.2046 *m/z* ion is an MS contaminant that enters MS/MS because it falls within the isolation window for 394.1357 *m/z* targeted MS/MS.

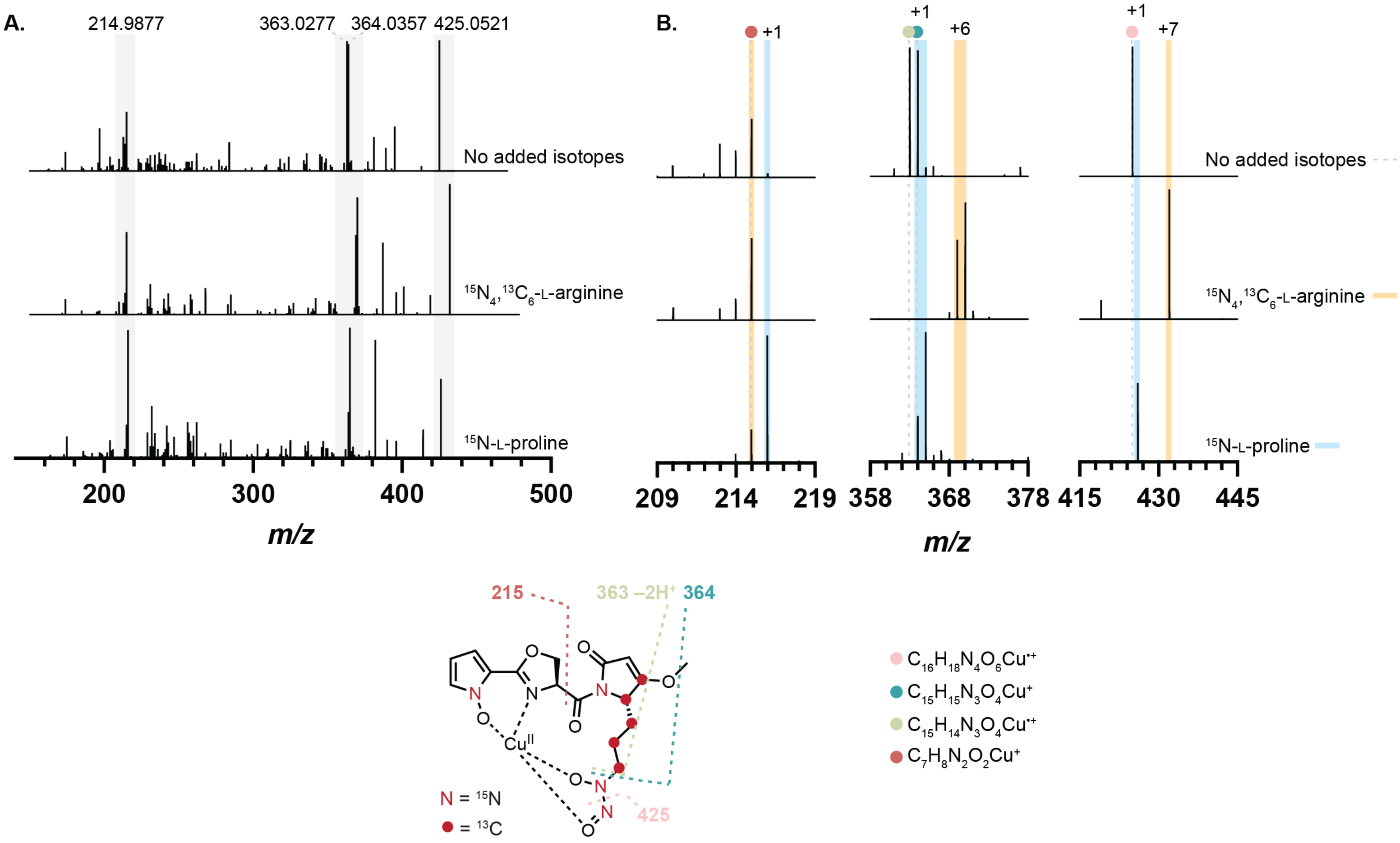


**Figure S5: LC–MS/MS of chalkophomycin ion for stable isotope feeding experiments. A.** MS/MS spectra of chalkophomycin when no stable isotopes, $^{15}$N-L-proline, or $^{15}N_4$, $^{13}C_6$-L-arginine were added to *S.* sp. Root63 cultures. The *m/z* ions listed above the spectra correspond to the values observed for the no added isotopes condition. The grey regions highlight features that are examined more closely in panel B. **B.** A +1 *m/z* shift is observed for all highlighted ions in the $^{15}$N-L-proline supplemented sample, all of which include the *N*-hydroxypyrrole moiety. No mass shift is observed for the 215 *m/z* ion in the $^{15}N_4$, $^{13}C_6$-L-arginine supplemented sample, consistent with all isotopes being incorporated to the right half of the molecule. The +6 *m/z* shift for both 363 and 364 *m/z* ions are consistent with enrichment of five $^{13}$C atoms and one $^{15}$N atom. Combined with the +7 *m/z* shift in the 425 *m/z* ion, this further localizes arginine-derived atoms to the diazeniumdiolate portion of chalkophomycin. Data was acquired on a ThermoFisher Orbitrap IQ-X with the following MS/MS parameters: HCD@25.

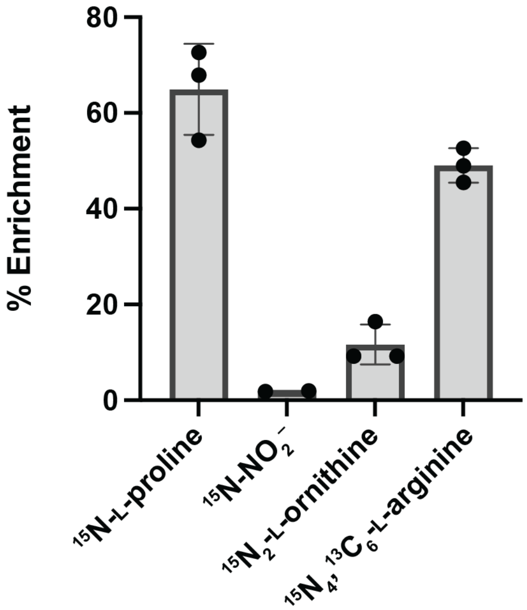


**Figure S6: Percent enrichment observed in stable isotope feeding experiments to determine chalkophomycin biosynthetic precursors.** Individual replicates are displayed as individual data points and error bars represent standard deviation.

**Table S5: Individual replicate and average values for percent enrichment observed in stable isotope feeding experiments.**

| | % Enrichment | | | | |
|---|---|---|---|---|---|
| **Substrate** | Replicate 1 | Replicate 2 | Replicate 3 | Average | Standard Deviation |
| $^{15}$N-L-proline | 72.63 | 54.28 | 67.88 | 63.93 | 9.52 |
| $^{15}$N-sodium nitrite | 1.94 | 1.82 | – | 1.88 | 0.08 |
| $^{15}N_2$-L-ornithine | 9.26 | 9.23 | 16.46 | 11.65 | 4.16 |
| $^{15}N_4$,$^{13}C_6$-L-arginine | 52.64 | 48.96 | 45.48 | 49.03 | 3.58 |

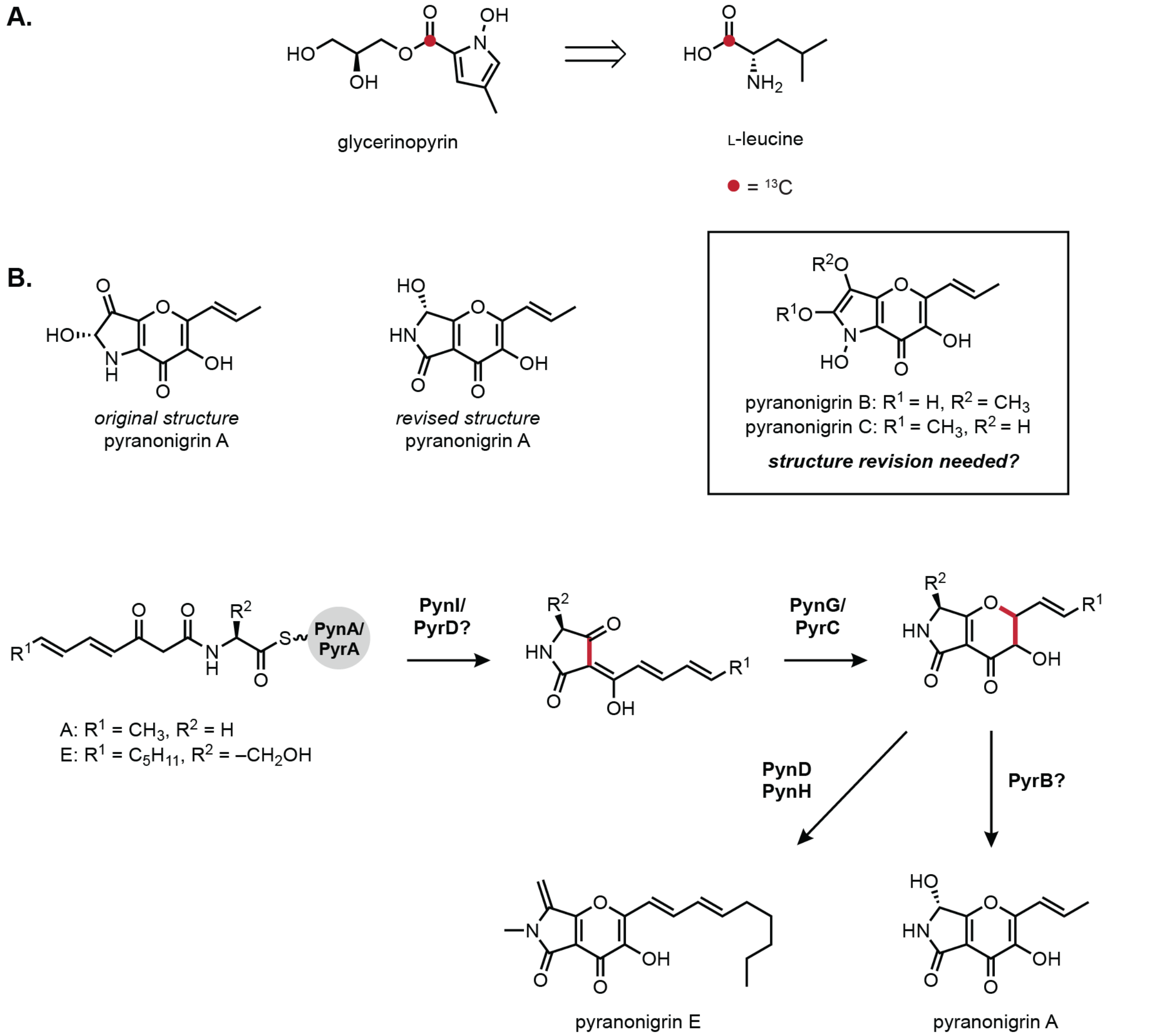


**Figure S7: The *N*-hydroxypyrrole functional group likely has several biosynthetic origins. A.** Glycerinopyrin biosynthesis is believed to originate from L-leucine according to stable isotope feeding experiments.[7] **B.** Genetic deletions, *in vitro* biochemistry, and stable isotope labeling support the pyranonigrin core scaffold arising from acetyl-CoA, malonyl-CoA, and an amino acid.[8–10] We note that the originally proposed structure of pyranonigrin A was later revised. The structure of pyranonigrin B and C, which were inferred from the originally structure of pyranonigrin A, has not yet been revised.

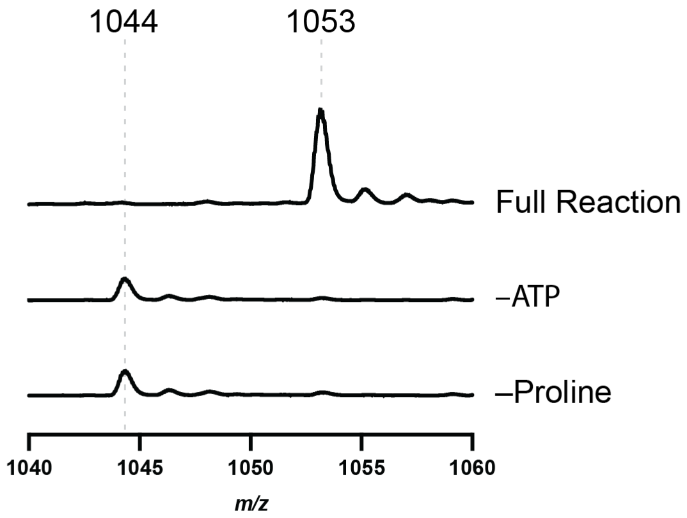


**Figure S8. ChmJ adenylates proline for loading onto ChmI.** Whole protein LC–MS of *in vitro* reaction of ChmI and ChmJ and controls without ATP and L-proline. Expected *m/z*: $[\text{ChmI+ppant+11H}]^{11+}$ = 1044.4167, $[\text{ChmI+ppant+proline+11H}]^{11+}$ = 1053.2397.

```
ChmG/1-396      1 -MHATDT---PHTEPLLDRVRRTAEELRLRSAEVDRGTLTPHDNYRLVREAGLLTLLVPREHG--GAG-LSYLD-  67
ChmH/1-384      1 MTPGSTR---TDIEDTVAHAARLA-----ESALADADGADFRTRWKGLAARGVWDAPAPDS----GAGRLGPVTG  63
Bmp3/1-380      1 MNFEWT----HEQAELFEHALRFGKE---LSAPLQEDNGFPRDNWNALGDFGYFGLPIPEKYAKDGSG-FDILT-  66
PltE/1-380      1 MDFNYD----DTQKKHAAMIAQVCAE-QLAACGNEHSRYFTARQWAICGEAGLLGLSIPREYG--GQG-LGALS-  66
RedW/1-391      1 MNFDFDAGFDTETRELRDMVVRFARR-ELDSSGRFDDAEDFRRRWLLAGKQGLTGTTVPGEYG--GSG-LDAVS-  70

ChmG/1-396     68 YTKVLETLAAGDGATALGFNMHNVAIGSLCETGGTGLP-PGASRFRDWVFGEVVEHSRMFASATSETGSGAKLRR 141
ChmH/1-384     64 AVATVEGIGRAGTAAGLCYAMASQRFGIQFPL----LSVLGEDGPRR--LGDTGSGDVLLCHALTEEGGGSDPLS 132
Bmp3/1-380     67 TIKIIEGLGQSCTDTGLLFAGAAHTFACSMPI----LE-HGSETLKHQLLPDLATGRKIAANAISEASAGSDISN 136
PltE/1-380     67 TAIAMHAFGLGCTDMGLVFAAAAHQFACAMPI----VE-FATAETKRDVLPKLASGEFIGSNAITEPEAGSDSSN 136
RedW/1-391     71 AAATMEALGYGCADTGFAFSVAAHLFAAVMPI----VE-FGTGEQRAAWLPALCSGERIAAHAITEPEAGSDALH 140

ChmG/1-396    142 IQATYRTSGDG-YILKGTKAFVSLAGIADHYVVAAREDGTSEADEVSHFVVSRDDPGVAFSGVWDGAALRGTQTA 215
ChmH/1-384    133 MSTRAELQEDGGYLLTGTKSFVTAAPVADVALVFARTAAERSPFALSAFLVDLRSPGVEQSAPFPKTALTEVPMG 207
Bmp3/1-380    137 LATTAQKEGDY-YVLNGGKSYVTNGSIADYYVVYATTNKKHGYLGQTAFVVPRNTPGISVGNDYHKLGLRSAPLN 210
PltE/1-380    137 LKSRAWPQADGSYRLDGHKSFAGNAPIADIFVTYATTQPEYGALGVSGFIVHRSSAGLRVSEPLDKVCLRSCPAG 211
RedW/1-391    141 LRTRARPVDDG-HVLSGSKCFITNAPVADVFVVQAATDPRGGFFGLTTFLVEASTPGLTVGRPYDKVGLRGSPTA 214

                                                 *
ChmG/1-396    216 TMTMNEVEVPRHRLFLGVEGLSLFKLVREPHWMVSGYMGAYLGIAESILRLMVRLLKDNE--GRR--SSPVVQAE 286
ChmH/1-384    208 SITFHGVRLGPDRMV-GEEGAGLGLLTLTTAWERALLLSYALGPMRRVLDRTVEWSATRQHFGRRMGASHIVAAR 281
Bmp3/1-380    211 QVFFDNCTIHKDYAL-GREGQGARIFAASMDWERCCLFAIFVGAMQRDLNQCIEYANTRMQGDKTISRFQAVSHR 284
PltE/1-380    212 EVFFDDCRVPEVNRL-GEEGQGRQVFQSSMGWERACLFAAFLGMMERQLEQTIEHARTRRQFGKPIGDNQAVSHR 285
RedW/1-391    215 DVHFDDCYVPAGAVL-GAEGSGASIFSSSMKWERTCLFAAYLGAMRRVLESTVDHVRDREQFGSPIGGFQAVSHR 288

ChmG/1-396    287 VGRLAVELRAARALVHSAARQVDEGRGSLEANTAVHAAKYCVGELAPRLALGAARICGSGALRTSGPLERLLREA 361
ChmH/1-384    282 VADMALALYRSRELVYRMAARLDAGERPRQLASDAALTKISVAEDYLLFSQQAARLGGVRSFIEDSGLTADLTSP 356
Bmp3/1-380    285 IADMGVRLESARLMLYYAAWQKSQDV---DNTKAVAMSKLAISEAFVQSGIDSIRVHGALGYLDEGRVNNSIKDA 356
PltE/1-380    286 IAQMKLRLESARLLLFRACWGMDQGD---PGQLNIALSKLAISEGALASSIDAVRIFGGRGCLESFGIEAMLRDS 357
RedW/1-391    289 IVDMLGRYEGARLLLYRAARSLSDGT---ADEVGPALAKIAVSEAAVQLGLDAVQLRGGLGIMD-GEAETLLRDA 359

ChmG/1-396    362 AFCAVMPAKPDECLEYVGKSTLGFNLFDAQNFDW*                                         396
ChmH/1-384    357 MAASTYAGPNDLLRITVARE---LGL----PVEN*                                         384
Bmp3/1-380    357 LGSVLFSGTSDIQRELICNR---LGLL--------                                         380
PltE/1-380    358 IGTTIFSGTSDMQHEIIARE---LKL---------                                         380
RedW/1-391    360 LPARIFSGTNEIQKNNVARA---LGLGRRRPAARR                                         391
```

**Figure S9: Multiple Sequence Alignment of ChmG and ChmH with a selection of known proline oxidases.** Sequences were aligned using MUSCLE algorithm in Geneious[11] with default parameters and colored using the Clustal color scheme in Jalview[12]. The catalytic Glu required for characterized proline oxidases is indicated by an asterisk (*).

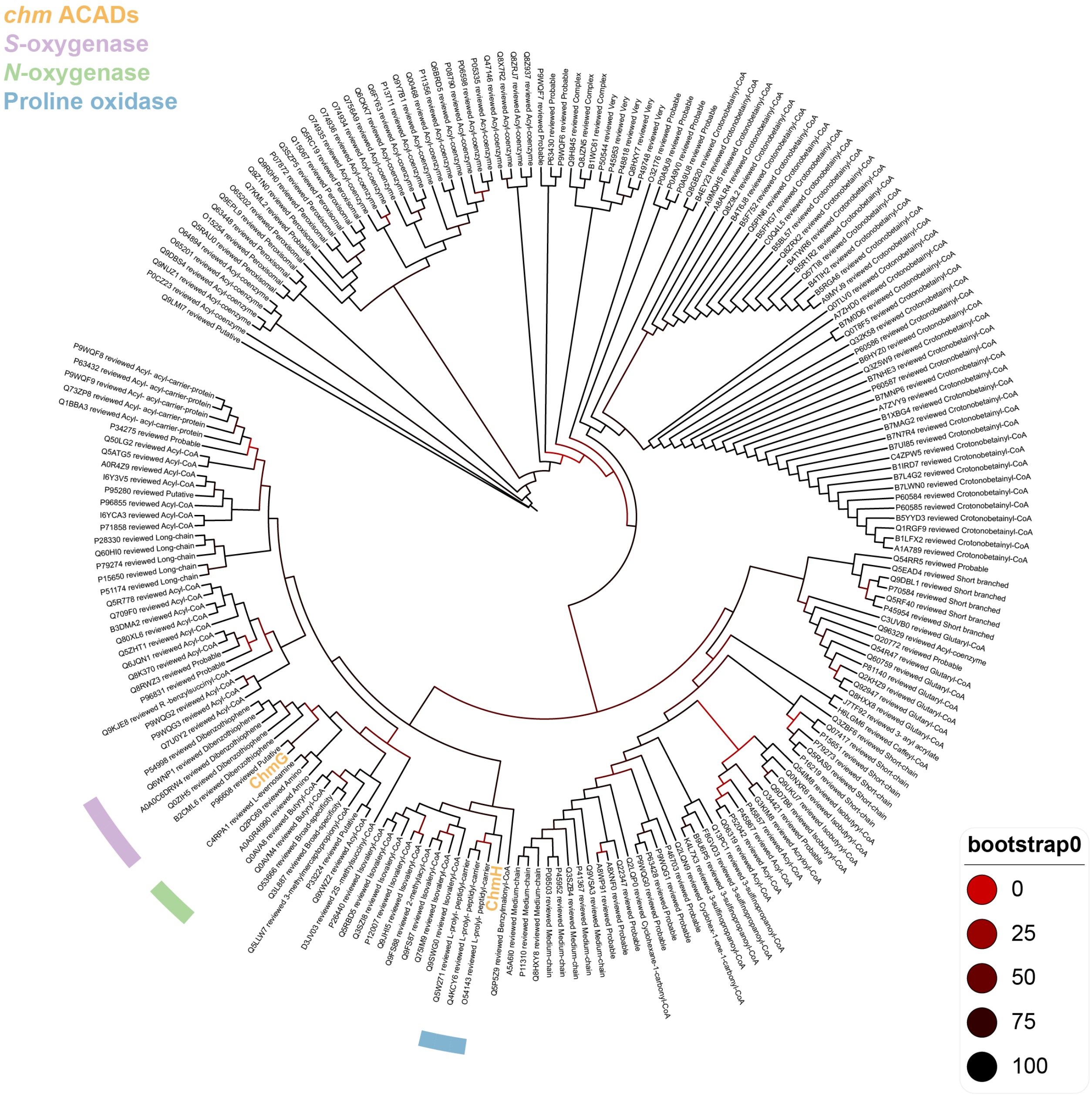


**Figure S10: Phylogenetic tree provides insight into ChmG and ChmH function.** Maximum likelihood phylogenetic tree of the reviewed sequences (n = 211) of InterPro family IPR006091 (Acyl-CoA oxidase/dehydrogenase, middle domain) indicates ChmG is most related to heteroatom oxygenases and ChmH to proline oxidases.

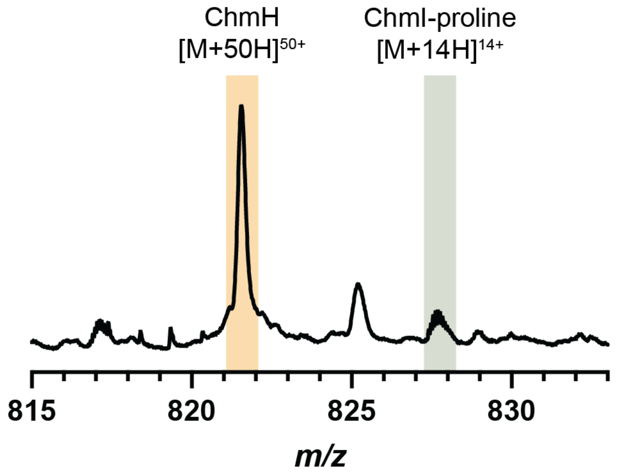


**Figure S11: *Lfl*ChmH is inactive towards ChmI–proline.** The +50 charge state of *Lfl*ChmH (Calc'd: 821.5573 *m/z*) and +14 charge state of ChmI-proline (Calc'd: 827.2707 *m/z*) are shown to provide greater resolution between the *Lfl*ChmH and ChmI-proline analytes and demonstrate no change to the ChmI-proline substrate mass feature.

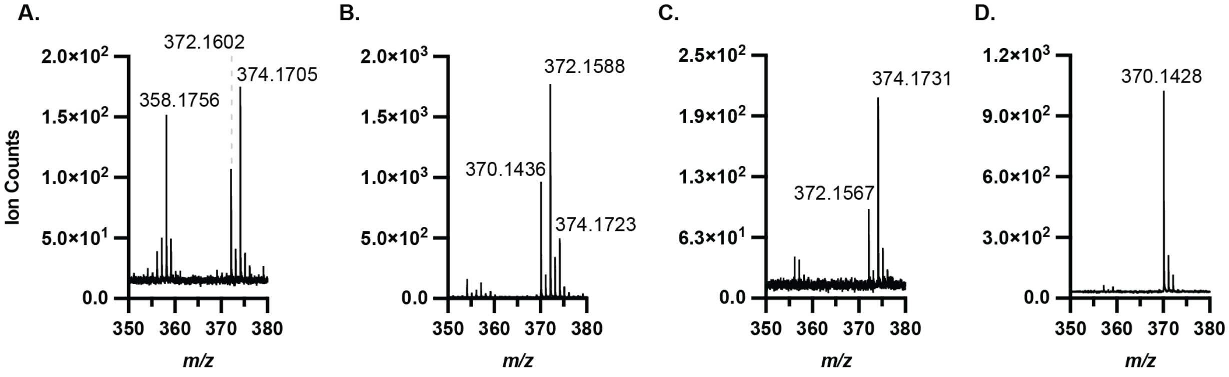


**Figure S12: FADH$_2$ availability influences *Lfl*ChmG product ratios.** MS/MS fragmentation of *in vitro* reaction of ChmIJ and *Lfl*ChmG products shows that **A.** in the absence of NADPH the major product is ChmI-*N*-hydroxyproline (MS/MS expected mass: 374.1744 *m/z*) while **B.** addition of a flavin reductase results in the major product being ChmI-*N*-hydroxydehyroproline (MS/MS expected mass: 372.1588 *m/z*) along with the production of a new mass consistent with ChmI-*N*-hydroxypyrrole (MS/MS expected mass: 370.1431 *m/z*). **C.** When superoxide dismutase and catalase are added to the ChmIJ + *Lfl*ChmG reaction, the product ratio is shifted towards ChmI-*N*-hydroxyproline, although both products are still formed. **D.** Addition of superoxide dismutase and catalase to the ChmIJ + *Lfl*ChmG and *Lfl*ChmH reaction does not influence formation of ChmI-*N*-hydroxypyrrole.

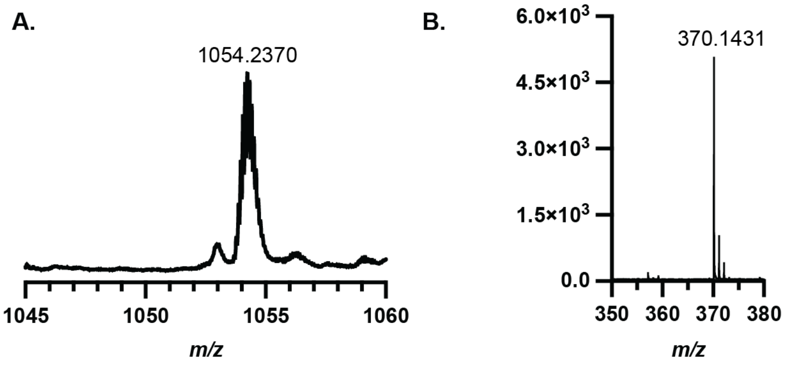


**Figure S13: Sequential addition of *Lfl*ChmH to ChmIJG reaction results in production of *N*-hydroxypyrrolyl-ChmI.** When *Lfl*ChmG is pre-incubated with ChmI-proline for 30 minutes prior to addition of *Lfl*ChmH, the same product (*N*-hydroxypyrrolyl-ChmI) can be observed by **A.** MS and **B.** MS/MS as found in reactions were *Lfl*ChmG and *Lfl*ChmH are added at the same time. This supports the hypothesis that the *N*-hydroxyproline and *N*-hydroxydehydroproline-ChmI thioester products generated by *Lfl*ChmG are both on pathway and can be oxidized by *Lfl*ChmH.

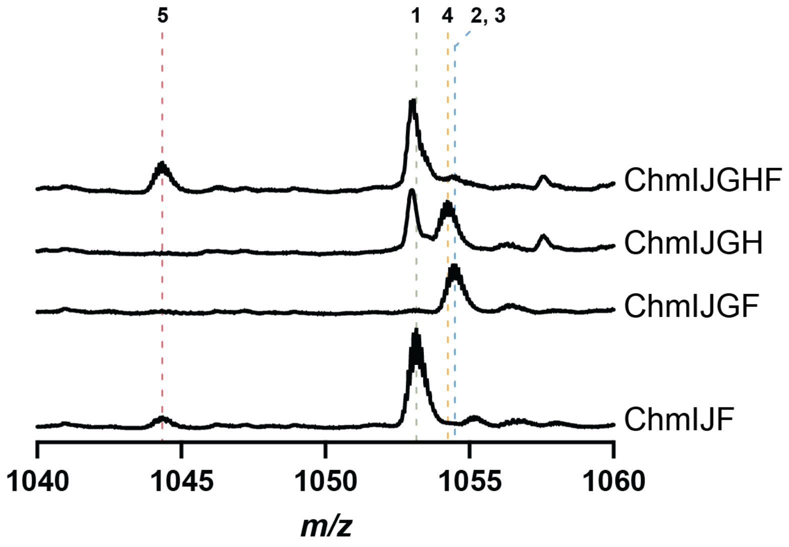


**Figure S14: ChmF hydrolyzes the *N*-hydroxypyrrole product from ChmI.** Whole protein LC–MS-based activity assay demonstrates that ChmF hydrolyzes *N*-hydroxypyrrole from ChmI, as evidenced by disappearance of **4** (*N*-hydroxypyrrolyl-ChmI) and appearance of **5** (holo-ChmI). This activity seems to be most abundant for the *N*-hydroxypyrrole; however, a small amount of substrate-free ChmI can be observed for the ChmIJF reaction, indicating ChmF can hydrolyze the prolyl thioester in low levels.

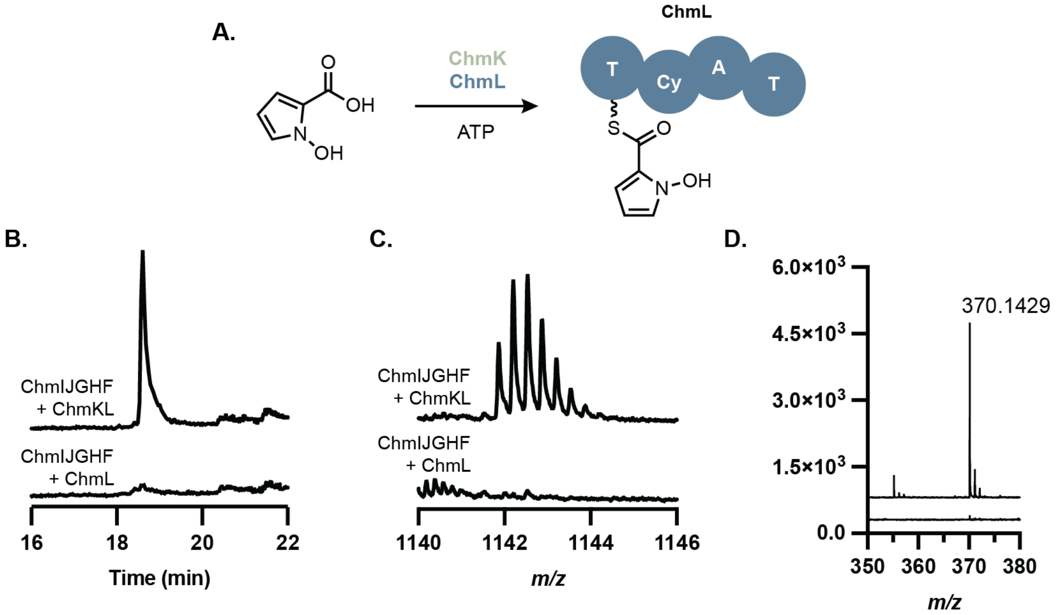


**Figure S15: ChmK adenylates N-hydroxypyrrole-2-carboxylic acid and transfers it onto the thiolation (T) domain of ChmL. A.** Scheme of ChmK activity. **B.** EIC for trypsin digest fragment of ChmL containing an *N*-hydroxypyrrolyl-ppant modification. Expected *m/z*: [A LVAGELDVDPAAIGPDEDLVTLGLHSMR+ppant+*N*-hydroxypyrrole+3H]$^{3+}$ = 1141.8789. **C.** Average mass spectrum extracted from the EIC in **B**. **D.** MS/MS of the mass feature presented in **B.** shows diagnostic ppant-*N*-hydroxypyrrole fragmentation. MS/MS expected mass: 370.1431 *m/z*.

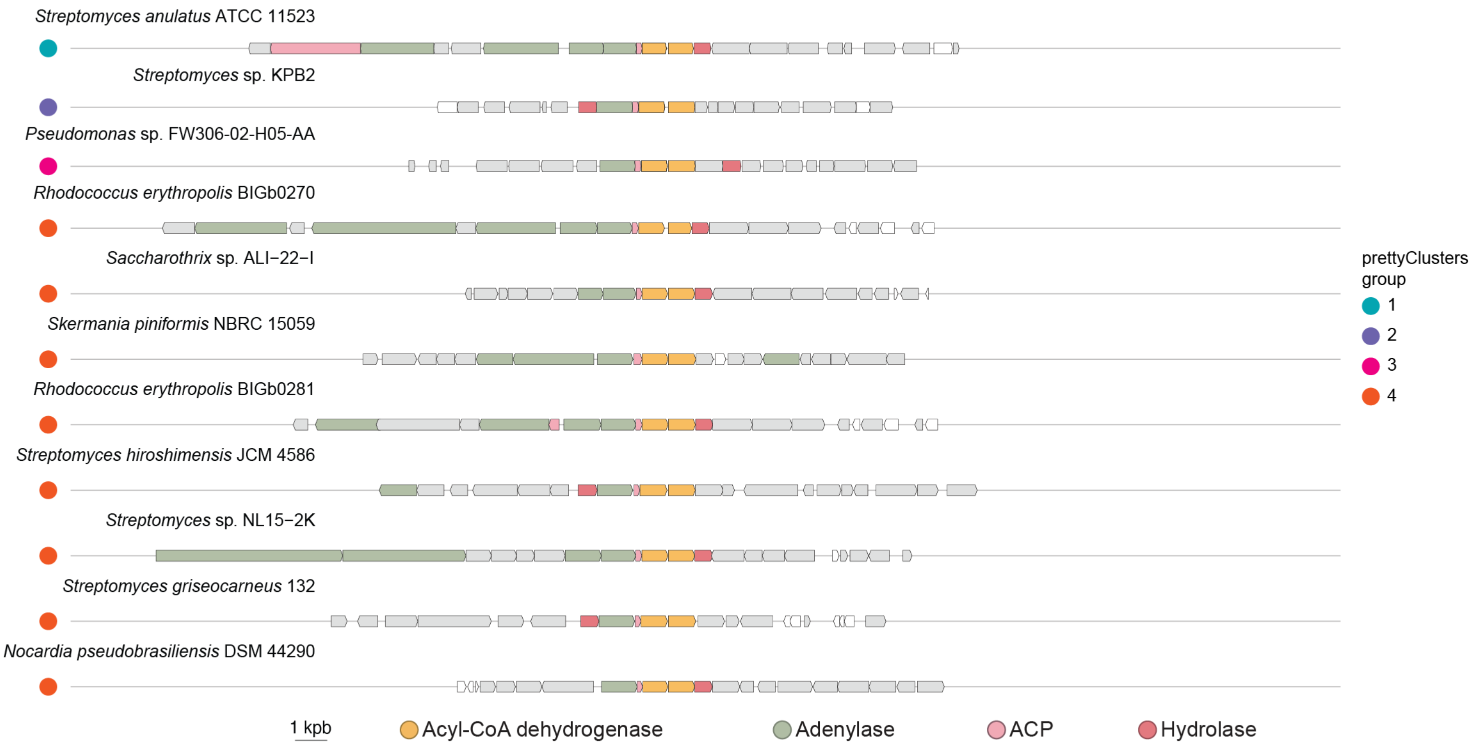


**Figure S16: There is variety within *chmIJGH* homolog-containing gene clusters.** Representative gene clusters are shown for each of the 4 cluster types categorized by prettyClusters13 to contain chmIJGH homologs. The fourth group had significant diversity, and as such shown here is a representative selection of 7 gene clusters. Group 1 (n = 10), Group 2 (n = 31), Group 3 (n = 20), Group 4 (n = 36).

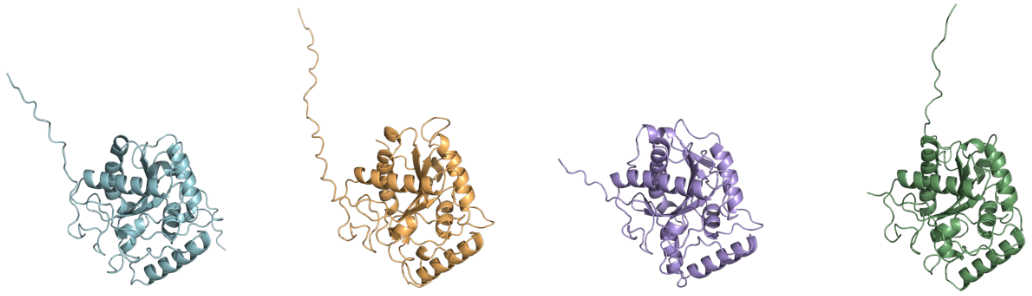

**Figure S17: Structural alignment of ChmN, AglA, GntA, and DcsA AlphaFold predictions.** Structures were generated using ColabFold and visualized in PyMOL using the “align” command. ChmN (blue), AglA (orange), GntA (purple), and DcsA (green) share a conserved tertiary structure.

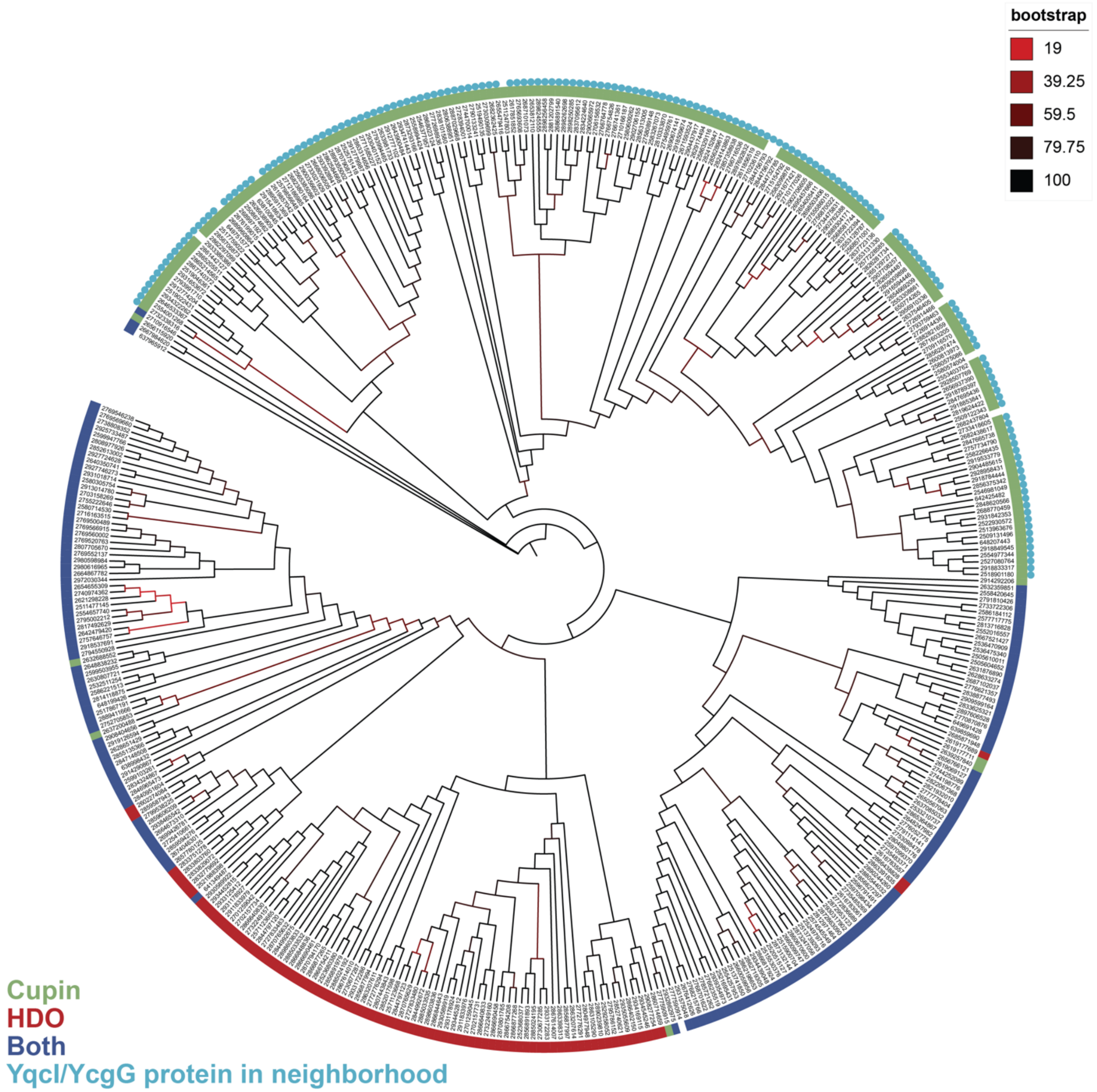


**Figure S18: Maximum likelihood phylogenetic tree shows conservation of cupin domain only SznF homologs with genes encoding for members of the YqcI/YcgG protein family.** Phylogenetic analysis of the same 426 protein sequences included in the Figure 1 SSN colored by conservation of the metal-binding residues in the HDO and/or cupin domains. prettyClusters was used to record the presence of genes annotated as members of pfam08892 (YqcI/YcgG family) within 10 genes distance of the *sznF* homologs.

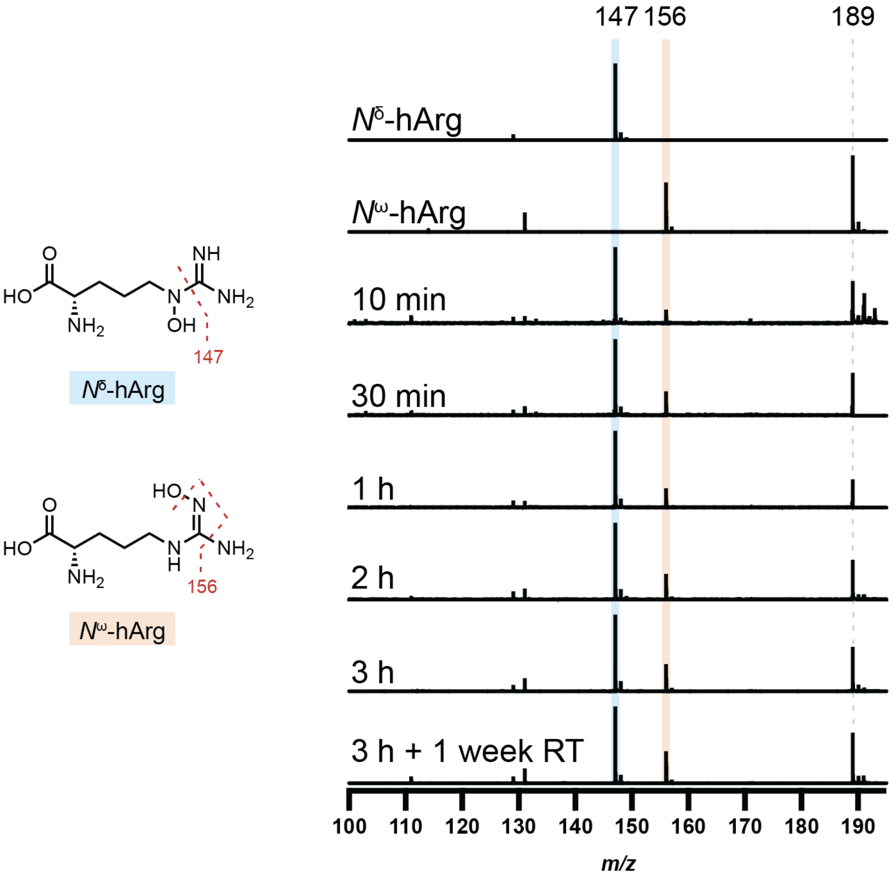


**Figure S19: LC–MS/MS analysis of ChmN time course.** MS/MS analysis of the 189.0993 *m/z* ion from ChmN *in vitro* assays, corresponding to hydroxylated arginine. Increasing counts of the 156 *m/z* fragment ion correspond to an increasing concentration of $N^{\omega}$-hydroxyarginine over time. Frag. = 175 V, CID@5

**Table S6: Ratios of 147:156 *m/z* fragment ions.**

| Time | Ratio *m/z* 147:156 |
|---|---|
| 10 min | 5.52:1 |
| 30 min | 3.17:1 |
| 1 h | 4.03:1 |
| 2 h | 3.05:1 |
| 3 h | 2.88:1 |
| 1 week at RT after quench | 2.40:1 |

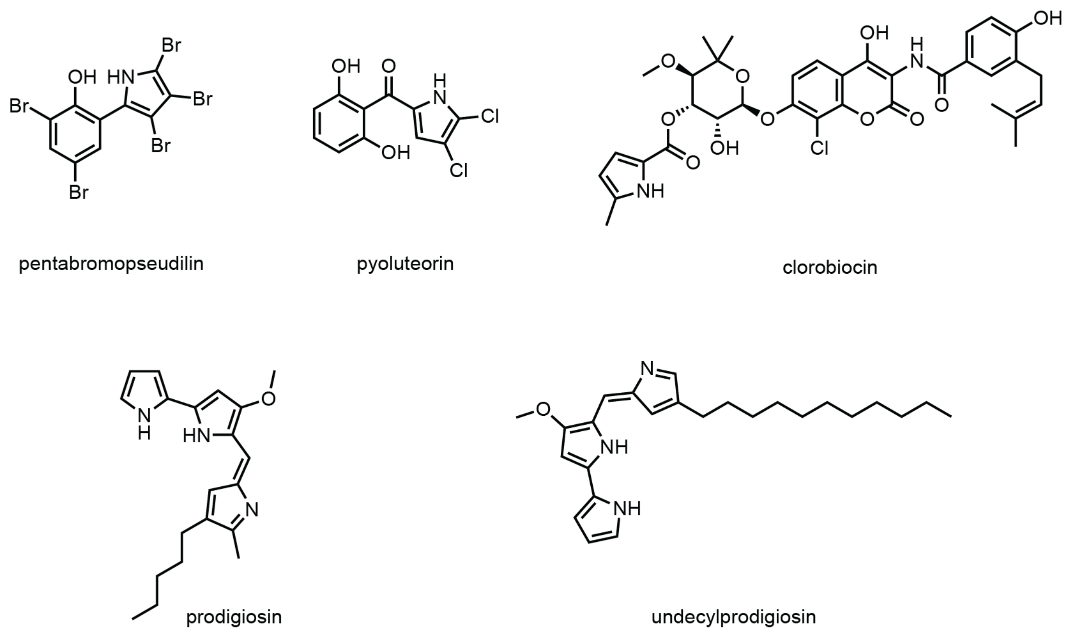


**Figure S20: Selection of pyrrole-containing natural products discussed in this manuscript.**

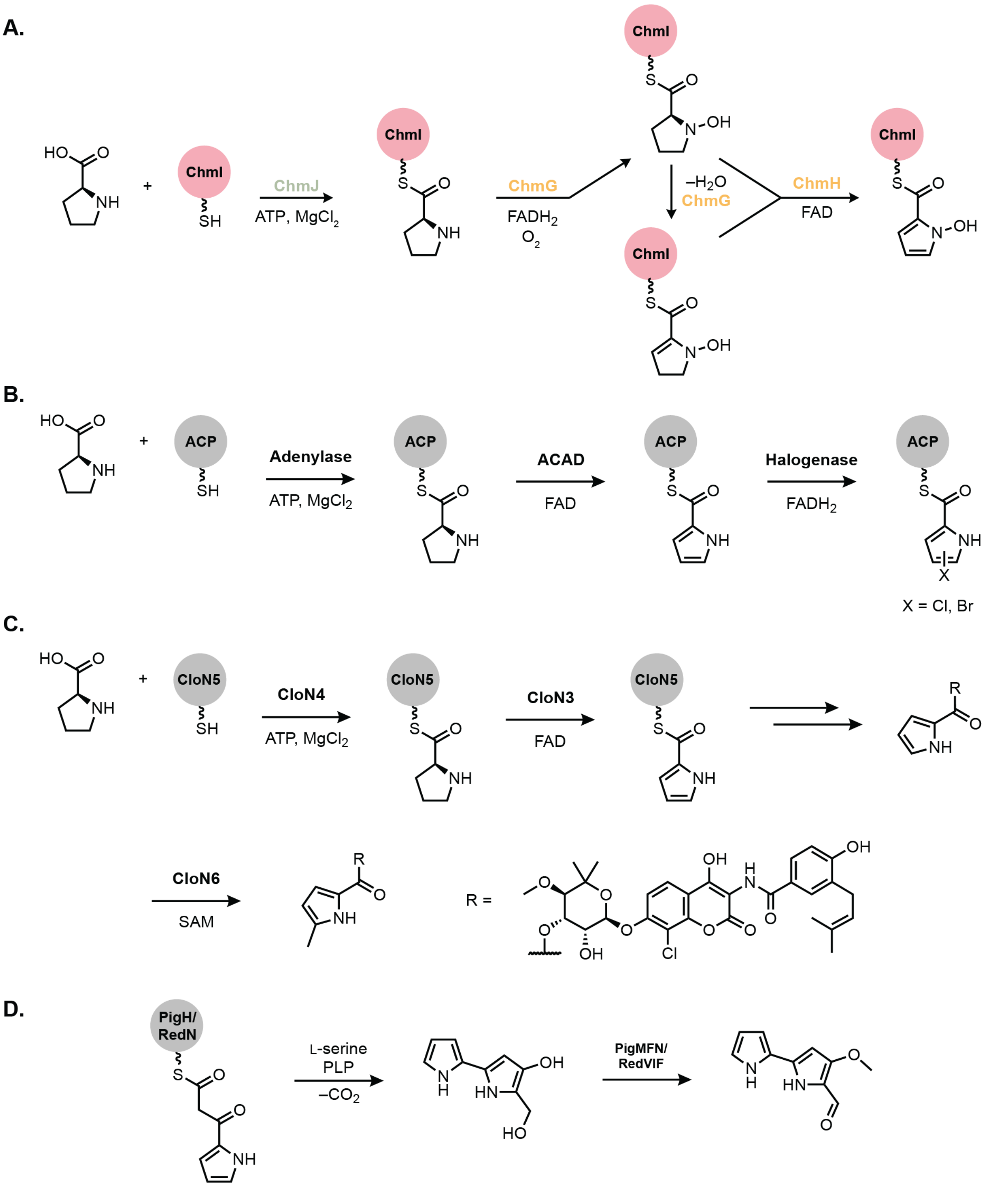


**Figure S21: Strategies for functionalized pyrrole biosynthesis. A.** *N*-hydroxypyrrole biosynthesis reported in this study. **B.** Halopyrrole (pyoluteorin, pentapromopseudilin) and **C.** methyl pyrrole (clorobiocin) biosynthesis follow the order of functionalization after proline oxidation. **D.** Methoxypyrrole (prodigiosin, undecylprodigiosin) biosynthesis utilizes a non-proline derived pathway.
